# Single-cell atlas reveals age-related cellular shifts underlying fibrosis in murine synovium

**DOI:** 10.64898/2026.04.17.719179

**Authors:** Olivier Bortolotti, Léa Marineche, Bienfait Abasi-Ali, Dany Séverac, Felicia Leccia, Christophe Duperray, Lucas Brugioti, Jacques Colinge, Christelle Bertrand-Gaday, Salwa Sebti, Florence Apparailly, Gabriel Courties

## Abstract

Aging is a major risk factor for joint disease, yet the impact of physiological aging on the synovium remains poorly defined. Here we generate a single-cell atlas of murine ankle synovium across age and identify the sublining stromal-myeloid niche as a major site of age-associated remodeling. Aging shifted fibroblast states toward oxidative stress and matrix-remodeling programs, accompanied by sublining collagen accumulation, reduced cellularity, and loss of THY1^+^ sublining fibroblasts. In parallel, resident synovial macrophages exhibited altered inflammatory and phagocytic responses together with a preferential decline in TIM4^+^VSIG4^−^ sublining macrophages, without overt local myeloid expansion despite systemic inflammaging. Macrophage depletion experiments further supported a link between sublining macrophages and extracellular matrix homeostasis. Together, these findings provide a reference framework for synovial aging and uncover niche-specific stromal and macrophage alterations associated with aging.

---

Aging is the primary risk factor for progressive musculoskeletal dysfunction, driving debilitating joint disorders such as osteoarthritis (OA) and rheumatoid arthritis (RA) that severely impact quality of life^1,2^. Within the joint, the synovium forms a specialized connective tissue that supports homeostasis by nourishing the avascular cartilage, lubricating the joint cavity and providing immune surveillance. It is anatomically compartmentalized into two distinct niches: the lining layer, which directly faces the joint space, and the sublining layer, a deeper vascularized zone^3^. Recent studies have revealed that these layers are maintained by highly specialized networks of synovial fibroblasts and tissue-resident macrophages (TRMs). Together, these stromal-immune circuits regulate extracellular matrix (ECM) turnover, clear debris, and maintain a strict protective barrier against aberrant inflammation^4^. Given the homeostatic functions of these resident populations, age-associated alterations within this compartment are likely to contribute to joint vulnerability. However, research into joint aging has largely centered on cartilage degeneration, where key hallmarks such as cellular senescence, altered ECM remodeling, and chronic inflammation have been extensively characterized^5–7^. In contrast, how synovial tissue changes during unperturbed aging remains poorly defined. To address this question, we generated a single-cell transcriptomic atlas of murine synovium during physiological aging. By integrating single-cell RNA sequencing (scRNA-seq) with bulk transcriptomics, flow cytometry, and histological analyses, we define the cellular landscape of young and aged synovial tissue and identify the synovial sublining compartment as a major site of age-associated remodeling. This resource provides a framework for understanding how aging reshapes stromal-immune organization within the joint. It offers new insight into the cellular basis of age-related joint vulnerability.

## Results

### Single-cell transcriptome of young and aged mouse synovium

To map cellular and molecular changes in the synovial joint during unperturbed aging, we performed scRNA-seq on synovial tissue microdissected from ankle joints of young (2.5-month-old) and aged (24-month-old) C57BL/6J mice (Fig. 1a; Extended Data Fig. 1a, b). Following enzymatic dissociation, viable single cells were isolated by Fluorescence-Activated Cell Sorting (FACS) and processed using the 10x Genomics platform. We profiled 6,362 cells (young) and 7,058 cells (aged) passing stringent quality-control filters, including removal of low-quality cells and doublets (Extended Data Fig. 1c-e). Unsupervised clustering identified 17 major cell clusters spanning stromal, endothelial and hematopoietic compartments (Fig. 1b; Extended Data Fig. 2a; Supplementary Table 1). *Ptprc* (encoding CD45) and the stromal marker *Dcn* (decorin) delineated immune and stromal regions of the Uniform Manifold Approximation and Projection (UMAP), and canonical markers enabled annotation of fibroblast subsets, periarticular stromal and immune populations (Fig. 1c-e and Extended Data Fig. 2a,b). Based on high expression of *Pdgfra* and *Ly6a*, the stromal compartment comprised six fibroblast clusters (*Prg4*, *Myoc*, *Ugdh*, *Cxcl12*, *Col3a1*, and *Ccl11* fibroblasts), as well as *Ptn*^+^ osteoblast-like cells (*Runx2*, *Alpl*), *Angptl7*^+^ tenocyte-like cells (*Scx*, *Fmod*), and *Fabp4^+^* endothelial cells (*Pecam1*, *Cdh5*). We also detected a small *Mylpf*^+^ cluster (*Actn3*, *Tnnt3*) likely reflecting adjacent muscle captured during tissue isolation. Within the hematopoietic compartment, we identified three macrophage subtypes (*Cd74*, *Pf4*, and *Vsig4* macrophages), alongside monocytes (*Plac8*, *Ly6c2*), neutrophils (*Csf3r*, *Retnlg*), T lymphocytes (*Trbc2*, *Il2rb*), and mast cells (*Kit*, *Gata2*). Comparison of young and aged samples suggested age-associated remodeling of cell states in stromal and myeloid compartments, while canonical lineage markers remained broadly preserved across age (Fig. 1f and Extended Data Fig. 3). Differential expression analysis between young and aged conditions revealed 1,062 age-associated genes across clusters, with the largest number of DEGs observed in fibroblast and macrophage populations, highlighting the stromal-myeloid niche as a key site of age-associated remodeling (Fig. 1g; Supplementary Table 2). Together, these data provide a single-cell atlas of murine synovium in young and aged mice, establishing a framework for dissecting age-sensitive stromal and myeloid programs.

**Fig. 1:**
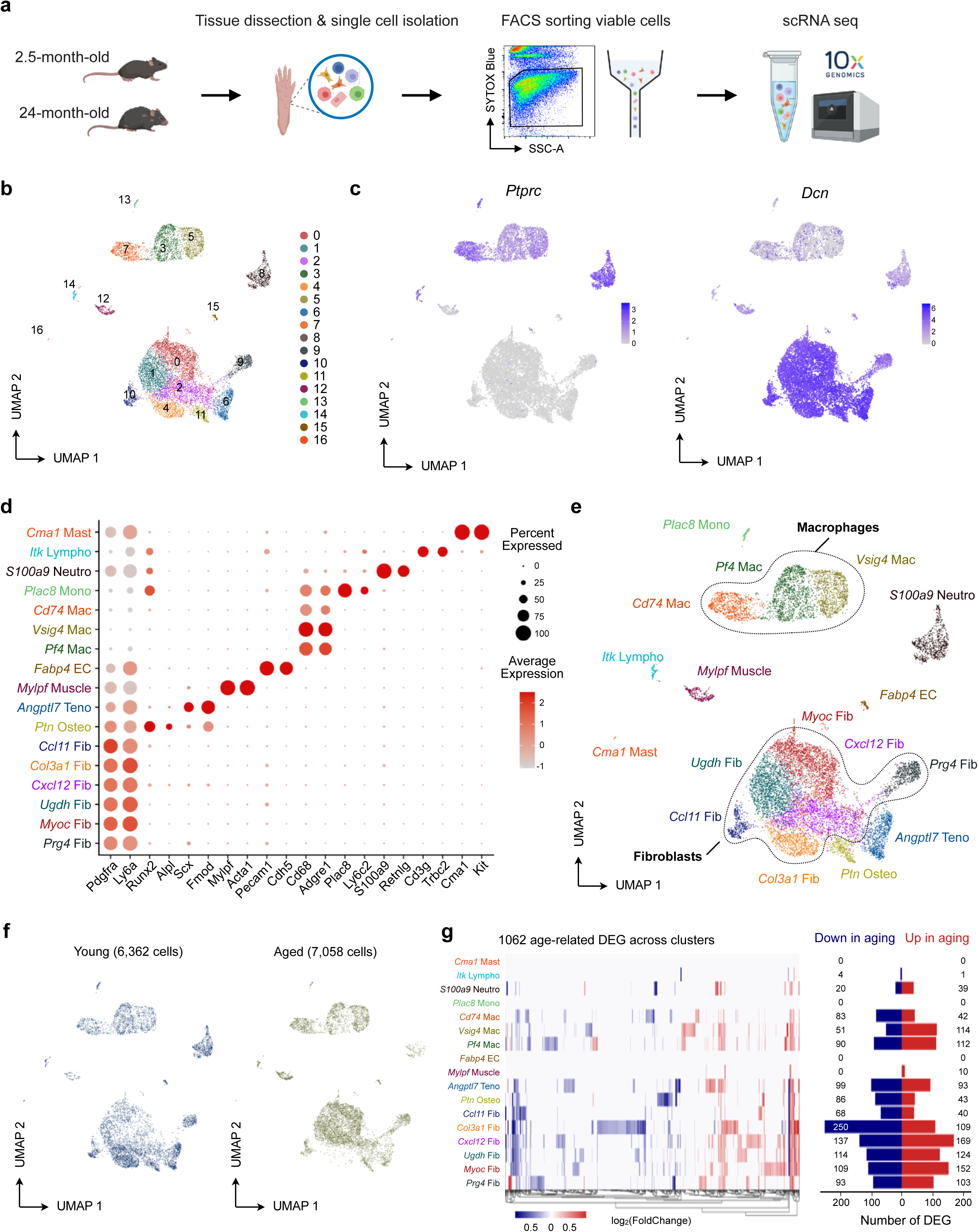
Single-cell transcriptional landscape of aging murine synovium. **a**, Workflow for synovial cell isolation, viable-cell sorting, and scRNA-seq from young and aged mice. **b**, Uniform manifold approximation and projection (UMAP) of the combined synovial cell dataset from young and aged mice, identifying 17 cell clusters. **c**, Feature plots showing expression of *Ptprc* and *Dcn*, delineating immune and stromal compartments. **d**, Dot plot of canonical marker gene expression across synovial cell clusters. Dot size indicates the percentage of cells expressing each gene, and color indicates average expression. **e**, Annotated UMAP of synovial cell populations based on differentially expressed genes (DEGs). **f**, UMAPs showing the distribution of synovial cell clusters in young and aged mice. **g**, Heatmap of age-associated DEGs across synovial cell clusters, with bar plots indicating the number of genes upregulated in aged mice (red) and downregulated in aged mice (blue).

### Aging reshapes synovial fibroblast heterogeneity toward a senescent-like state

Building on the synovial single-cell atlas, we focused on fibroblasts to define age-associated changes in stromal cell states. Fibroblasts segregated into *Prg4*⁺ lining fibroblasts (LFs) and *Thy1*⁺*Cd34*⁺ sublining fibroblasts (SLFs), the latter comprising five transcriptionally distinct subsets marked by *Myoc, Ugdh, Cxcl12, Col3a1* and *Ccl11* (Fig. 2a-c; Supplementary Table 3)^8,9^. These clusters were consistent with the murine synovial fibroblast taxonomy reported by Collins et al., with *Prg4*⁺ fibroblasts aligning with the FLS cluster and sublining fibroblasts distributed among F1-F6 states (Extended Data Fig. 4a)^10^. Functional annotation further supported biological specialization across subsets, with *Col3a1*⁺ and *Ccl11*⁺ fibroblasts enriched for matrix-supportive and angiogenesis-associated functions, whereas *Ugdh*⁺ fibroblasts were more strongly associated with stress- and inflammation-related pathways (Extended Data Fig. 4b). Comparison of young and aged samples revealed marked remodeling of the SLF compartment, with contraction of *Col3a1*⁺ and *Ccl11*⁺ SLFs and expansion of *Myoc*⁺ and *Ugdh*⁺ subsets (Fig. 2d). In parallel, Gene Ontology analysis indicated a shift away from tissue homeostasis programs including wound healing, connective tissue development, collagen organization and metabolic processes, toward pathways linked to inflammatory signaling, oxidative metabolism and cellular stress (Fig. 2e; Extended Data Fig. 4b). Intercellular communication analysis further showed that aging selectively rewired synovial signaling networks, with IL-6 emerging as a strengthened pathway in aged synovium and fibroblasts acting as both major senders and receivers (Fig. 2f and Extended Data Fig. 5a,b). SCENIC analysis likewise supported preservation of distinct fibroblast identities while highlighting subset-specific regulon programs, with the strongest enrichment of stress- and inflammation-associated regulons in the *Ugdh*⁺ state, including STAT3, NF-κB and AP-1 (Extended Data Fig. 5c)^11^. In line with this, the *Ugdh*⁺ cluster displayed the highest SenMayo score among fibroblast subsets (Fig. 2g)^12^. Together, these data indicate that aging remodels the synovial sublining fibroblast niche, shifting away from matrix-supportive homeostatic states toward a senescence-like profile.

**Fig. 2:**
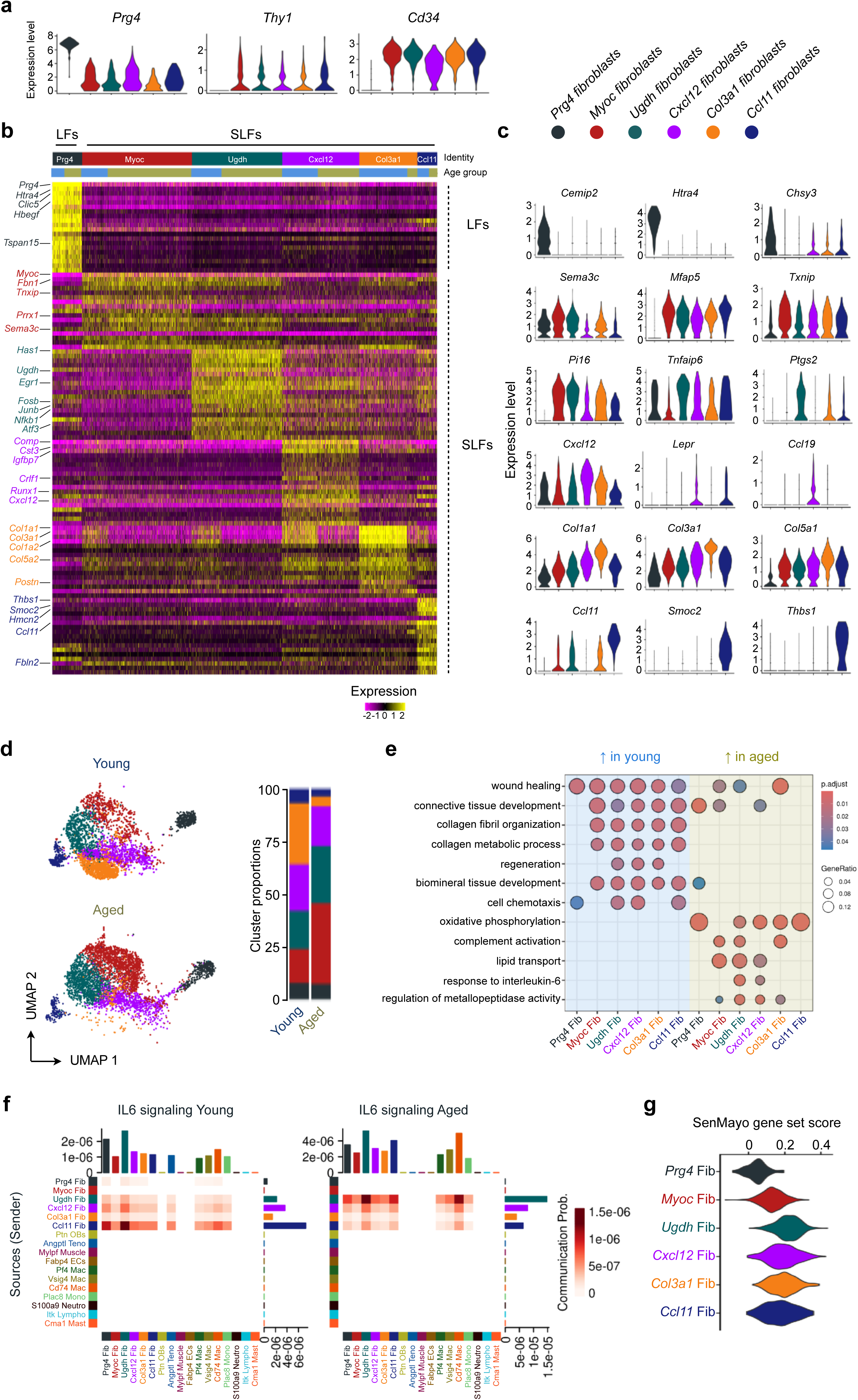
Heterogeneity and age-associated changes of synovial fibroblast populations. **a,** Violin plots showing expression of *Prg4*, *Thy1* and *Cd34* across fibroblast clusters. **b,** Heatmap of top 20 DEGs defining lining fibroblasts (LFs : *Prg4*⁺) and five sublining fibroblast populations (SLFs : *Myoc*⁺, *Ugdh*⁺, *Cxcl12*⁺, *Col3a1*⁺ and *Ccl11*⁺). **c,** Violin plots showing expression of representative marker genes across fibroblast subtypes. **d,** UMAPs showing fibroblast clusters in young and aged mice, with relative cluster proportions by age group. **e,** Gene Ontology enrichment analysis of genes upregulated in young versus aged fibroblast clusters. Dot size indicates gene ratio and color indicates adjusted *P* value. **f,** CellChat analysis of IL-6 signaling networks in young and aged synovium, showing inferred communication probabilities among cell populations. **g,** SenMayo module scores across fibroblast clusters.

### Aging remodels the synovium through fibrosis and preferential loss of THY1+ sublining fibroblasts

To assess whether these transcriptional changes were reflected at the tissue level, we performed histological analyses of young and aged synovium. Sirius Red staining revealed increased collagen deposition, predominantly within the aged sublining compartment, whereas immunohistochemistry showed higher fibroblast activation protein (FAP) expression, mainly within the intimal lining layer (Fig. 3a,b). This fibrotic phenotype coincided with reduced sublining cellularity, while lining thickness remained unchanged (Fig. 3c). Because THY1^+^ sublining fibroblasts are major stromal effectors in joint pathology, we next quantified synovial fibroblast subsets by flow cytometry^8,13–15^. Both ankle and knee joints contained fewer total PDGFRα^+^PDPN^+^ fibroblasts in aged mice, a change largely driven by preferential loss of the THY1^+^ sublining fibroblast population, whereas THY1^−^ lining fibroblasts were comparatively preserved (Fig. 3d,e). This reduction in sublining fibroblasts was also observed in female cohorts (Extended Data Fig. 6a,b). To further characterize age-associated transcriptional remodeling of the synovial fibroblast compartment, we performed bulk RNA-seq on FACS-sorted PDGFRα^+^PDPN^+^ fibroblasts. Principal component analysis separated young and aged fibroblasts along PC1, which accounted for 70% of the variance, and differential expression analysis revealed broad age-associated transcriptional changes (Extended Data Fig. 6c,d; Supplementary Table 4). Gene Set Enrichment Analysis corroborated the single-cell observations, showing reduced enrichment of extracellular matrix homeostasis pathways, including “collagen fibril organization”, together with increased enrichment of oxidative stress-related processes, including “hydrogen peroxide biosynthetic process” (Extended Data Fig. 6e). Aged fibroblasts upregulated oxidative stress-associated genes such as *Apoe, Txnip, Nox4* and *Duox1*, while also showing reduced expression of elasticity-associated genes (*Eln, Mfap4*) and increased expression of pro-fibrotic markers (*Tnc, Fap, Postn, Tgfb1*). CellROX-based flow cytometry confirmed significantly increased intracellular oxidative stress in aged PDGFRα^+^PDPN^+^ synovial fibroblasts supporting an oxidative stress-associated fibroblast state (Extended Data Fig. 6d-f). Thus, synovial aging is associated with fibrosis of the sublining compartment and preferential loss of THY1^+^ fibroblasts.

**Fig. 3:**
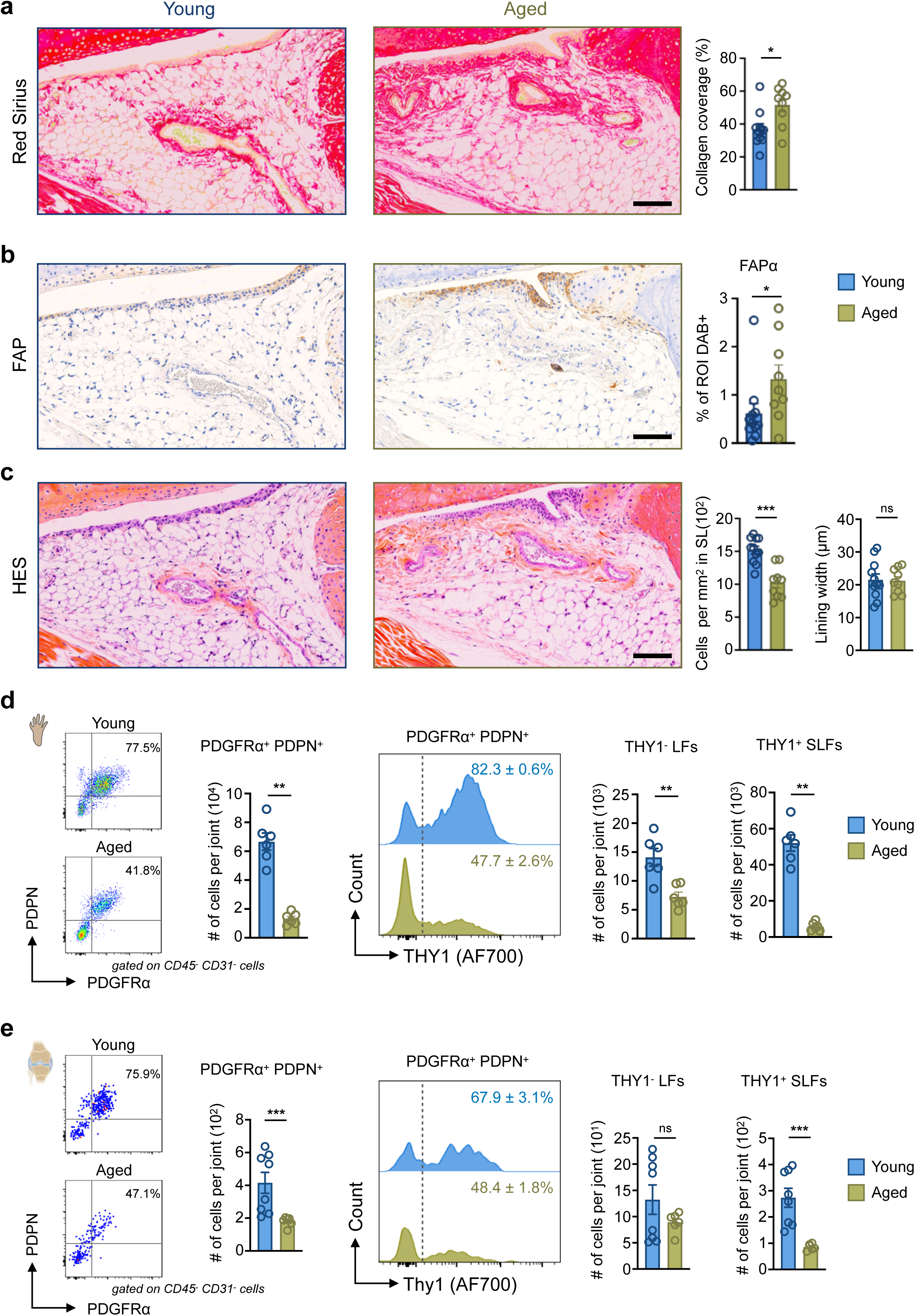
Sublining fibrosis and preferential loss of THY1+ fibroblasts characterize the aged synovium. **a,** Representative Sirius Red-stained sections of knee synovium from young and aged mice, with quantification of synovial collagen coverage. **b,** Immunohistochemistry for fibroblast activation protein (FAP) in knee synovium, with quantification of FAP-positive area. **c,** Hematoxylin–eosin–safran (HES) staining of knee synovium, with quantification of sublining cellularity and lining width (young n = 11 mice and aged n = 9 mice). Scale bars, 100 µm. **d,** Enumeration of CD45^−^CD31^−^PDGFRα⁺PDPN⁺THY1^−^ lining fibroblasts (LFs) and THY1^+^ sublining fibroblast subsets in ankle joints from young and aged mice by flow cytometry (n = 6 per group). **e,** Flow cytometric analysis of synovial fibroblast subsets from knee joints, shown as in d (young n = 8 mice and aged n = 6 mice). Data are mean ± s.e.m. Two-tailed Mann-Whitney *U* test. \**P* < 0.05, \*\**P* < 0.01, \*\*\**P* < 0.001; ns, not significant.

### Aging promotes systemic myeloid expansion without overt synovial infiltration

The histological and stromal changes observed in aged synovium indicated marked tissue remodeling, but they did not resemble the hyperplastic, leukocyte-rich phenotype typical of inflammatory synovitis^16,17^. Because aging is also associated with systemic inflammaging and myeloid-biased hematopoiesis, we next assessed systemic and local myeloid changes in young and aged mice^18,19^. In male mice, aging was associated with increased serum IL-6 levels together with significant expansion of circulating Ly6C^high^ monocytes and Ly6C^low^ monocytes (Fig. 4a,b). In contrast, flow cytometric analysis of synovial tissue revealed no significant increase in local F4/80^low^ Ly6C^high^ monocytes or Ly6G^+^ neutrophil numbers in aged joints (Fig. 4c). A similar uncoupling of systemic and local changes was observed in female cohorts, in which aging was also associated with increased serum IL-6 and elevated circulating myeloid cells, whereas tissue monocytes ansd neutrophils were not increased locally with age (Extended Data Fig. 7). Collectively, these findings indicate that physiological synovial aging develops in the setting of systemic inflammaging, but without overt accumulation of innate immune cells within the joint.

**Fig. 4:**
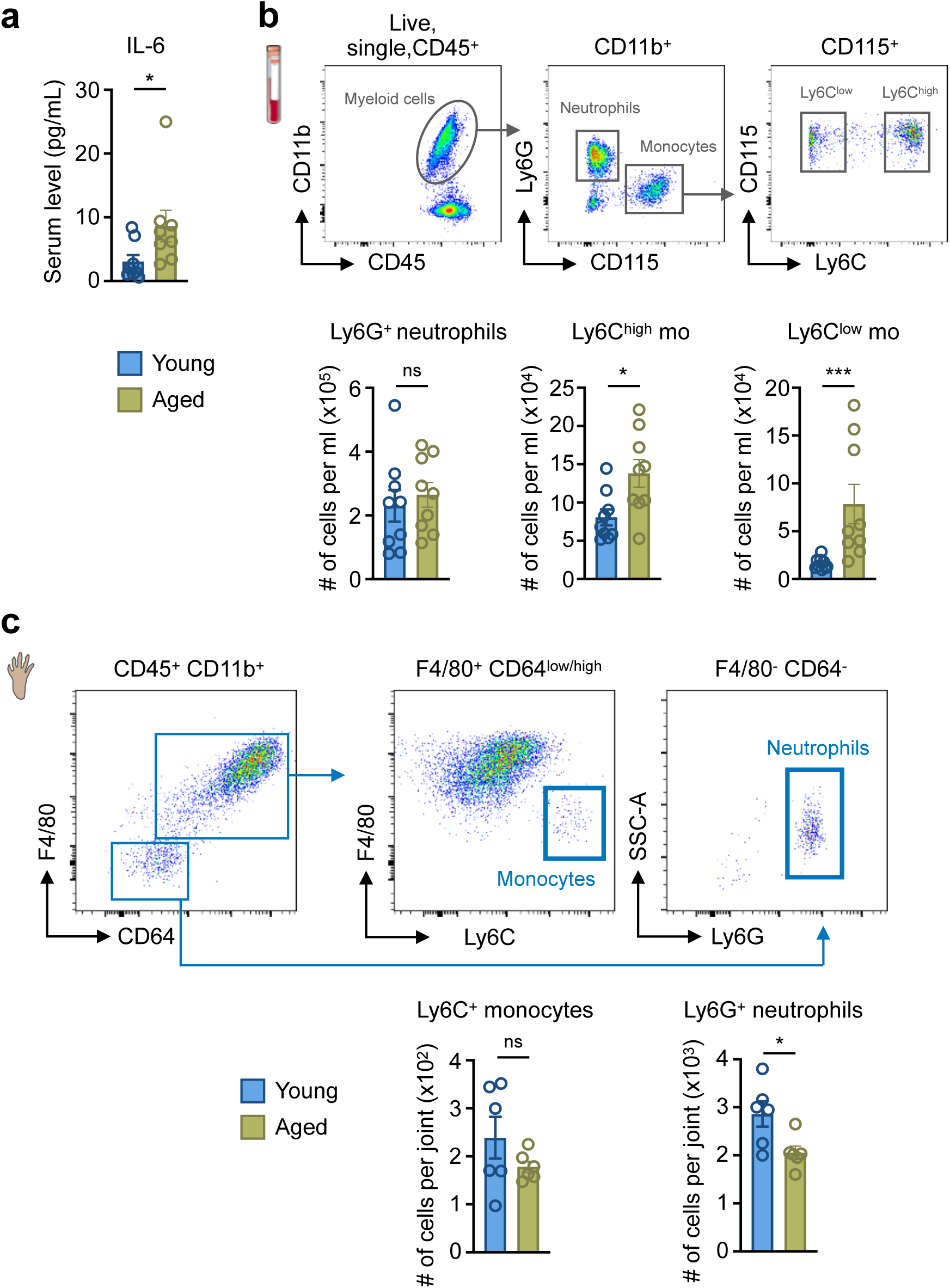
Systemic inflammaging occurs without overt synovial myeloid cell infiltration. **a**, Serum IL-6 levels in young and aged male mice (n = 8 per group). **b**, Representative flow cytometry gating strategy for peripheral blood myeloid cells. Bar graphs show the quantification of Ly6G^+^ neutrophils, Ly6C^high^ monocytes and Ly6C^low^ monocytes per ml of blood in young and aged male mice (n = 9 per group). **c**, Representative flow cytometry gating strategy for synovial tissue monocytes and neutrophils. Flow cytometry-based enumeration of Ly6C^+^ monocytes and Ly6G^+^ neutrophils per paw in young and aged male mice (n = 6 per group). Data are mean ± s.e.m. Two-tailed Mann-Whitney *U* test. \**P* < 0.05, \*\*\**P* < 0.001; ns, not significant.

### Resident synovial macrophages exhibit altered inflammatory responsiveness and phagocytic activity with age

Given the absence of overt myeloid cell recruitment in aged synovium, we next examined whether aging instead altered resident synovial macrophage populations. Single-cell RNA-seq identified three macrophage subsets distinguished by expression of *Pf4, Vsig4* and *Cd74* (Fig. 5a,b; Supplementary Table 5). In line with previous reports, the *Vsig4* cluster corresponded to lining-associated macrophages enriched for *Cx3cr1, Trem2* and *S100b*, whereas *Pf4* and *Cd74* defined two sublining populations with distinct transcriptional features (Extended Data Fig. 8a,b). In particular, the *Pf4* cluster aligned with interstitial macrophages enriched for *Aqp1, Mrc1* and *Retnla*, while the *Cd74* cluster matched MHCII-associated antigen-presenting macrophage subset^20–22^. Gene Ontology enrichment of differentially expressed genes indicated reduced representation of pathways linked to inflammatory responses, cellular responses to bacterial stimuli, leukocyte migration, wound healing and phagocytosis in aged mice across macrophage subsets (Fig. 5c). To test whether these transcriptional changes translated into altered macrophage activation and homeostatic functions, we evaluated inflammatory responsiveness and phagocytic activity in synovial macrophages isolated from young and aged mice. Synovial CD64⁺F4/80⁺ macrophages from aged joints produced significantly lower levels of IL-6 and TNF-α following ex vivo LPS stimulation than macrophages from young animals (Fig. 5d). In parallel, in vivo uptake of pHrodo-labeled *E. coli* particles showed a lower frequency of phagocytic macrophages in aged synovium (Fig. 5e). Together, these findings indicate that aging is associated with functional impairment of resident synovial macrophages, affecting both inflammatory responsiveness and clearance capacity.

**Fig. 5:**
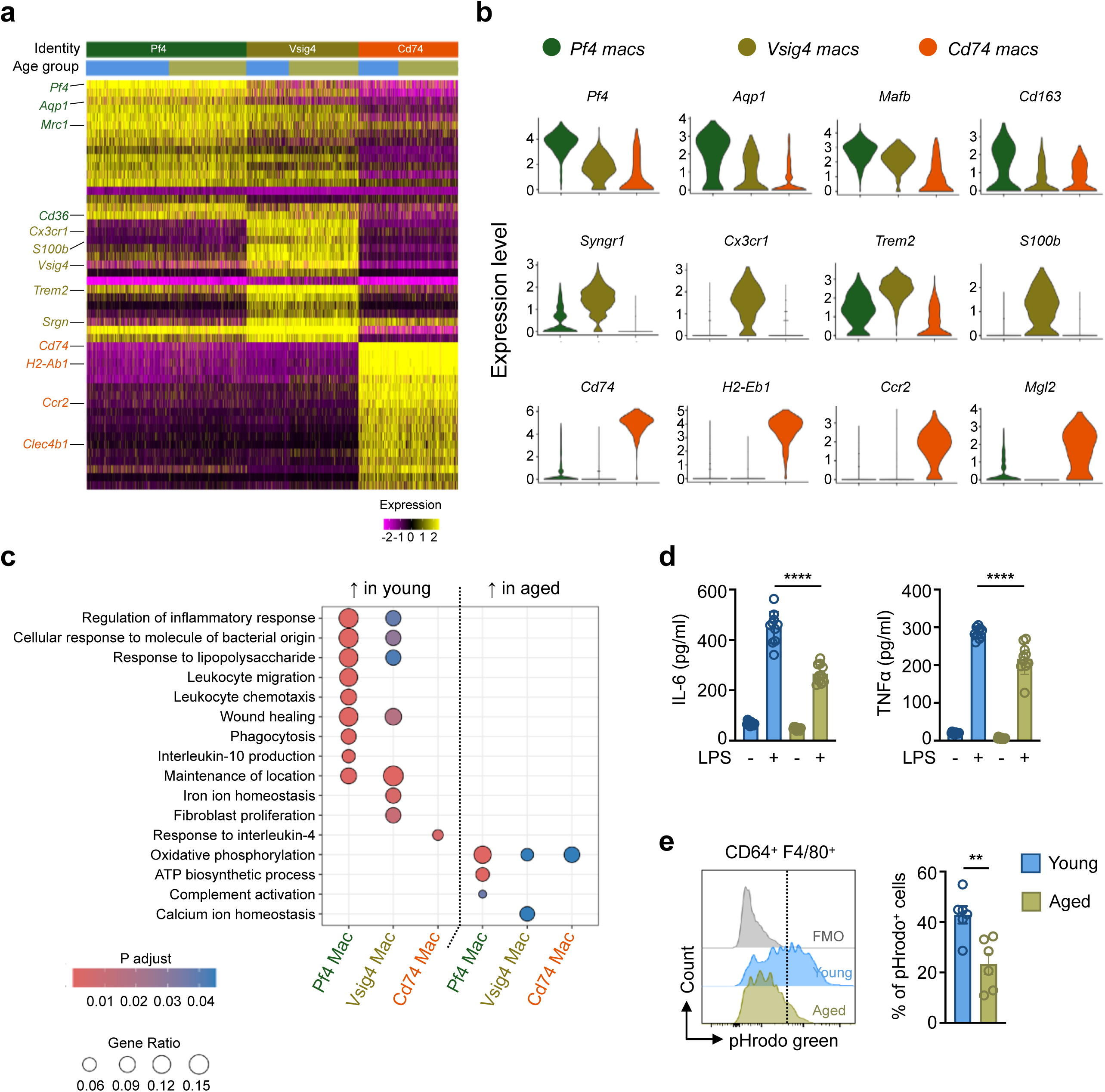
Synovial macrophages display age-associated functional impairments. **a**, Heatmap showing expression of top 20 genes across synovial macrophage clusters, identifying *Pf4*^+^, *Vsig4*^+^ and *Cd74*^+^ macrophage subsets. **b**, Violin plots depict expression of representative marker genes across the three macrophage subsets. **c**, Gene Ontology enrichment analysis of genes upregulated in young versus aged macrophages within each subset. Dot size indicates gene ratio and color indicates adjusted P value. **d**, IL-6 and TNF-α secretion by FACS-sorted synovial macrophages from young and aged mice following 24 h ex vivo stimulation with or without LPS (n = 10 mice per group; one-way ANOVA test). **e**, Phagocytosis assay showing representative pHrodo Green fluorescence histograms and frequency of pHrodo^+^ cells within CD64^+^F4/80^+^ synovial macrophages, 2 h after intra-articular injection of pHrodo Green E. coli BioParticles (n = 6 mice per group; two-tailed Mann-Whitney *U* test). FMO control is shown in grey. Data are mean ± s.e.m. \*\**P* < 0.01, \*\*\*\**P* < 0.0001.

### Aging preferentially reduces the TIM4^+^VSIG4^−^ sublining macrophage compartment

We next asked whether aging also affected specific resident macrophage niches within the joint. F4/80 immunohistochemistry showed a significant reduction in macrophages within the synovial subintima of aged joints, whereas no significant change was detected in the intimal lining layer (Fig. 6a). We next used TIM4 and VSIG4 cell surface markers to resolve resident synovial macrophage subsets identified in the single-cell atlas by flow cytometry. *Timd4* and *Vsig4* expression across macrophage clusters supported three corresponding populations: TIM4⁺VSIG4⁺ lining macrophages, TIM4⁺VSIG4⁻ sublining macrophages, and TIM4⁻ macrophages (Fig. 6b,c). Flow cytometric quantification demonstrated a marked reduction in the TIM4⁺VSIG4⁻ sublining macrophage population in aged mice across both ankle and knee joints, whereas TIM4⁺VSIG4⁺ lining macrophages showed a more limited age-associated reduction in both male and female mice (Fig. 6d,e and Extended Data Fig. 9a,b). Thus, the overall decline in synovial macrophages with age was driven predominantly by the loss of TIM4⁺VSIG4⁻ sublining macrophages. Together, these data indicate that the preferential loss of TIM4⁺VSIG4⁻ sublining macrophages is part of a broader age-associated defect of the synovial sublining compartment.

**Fig. 6:**
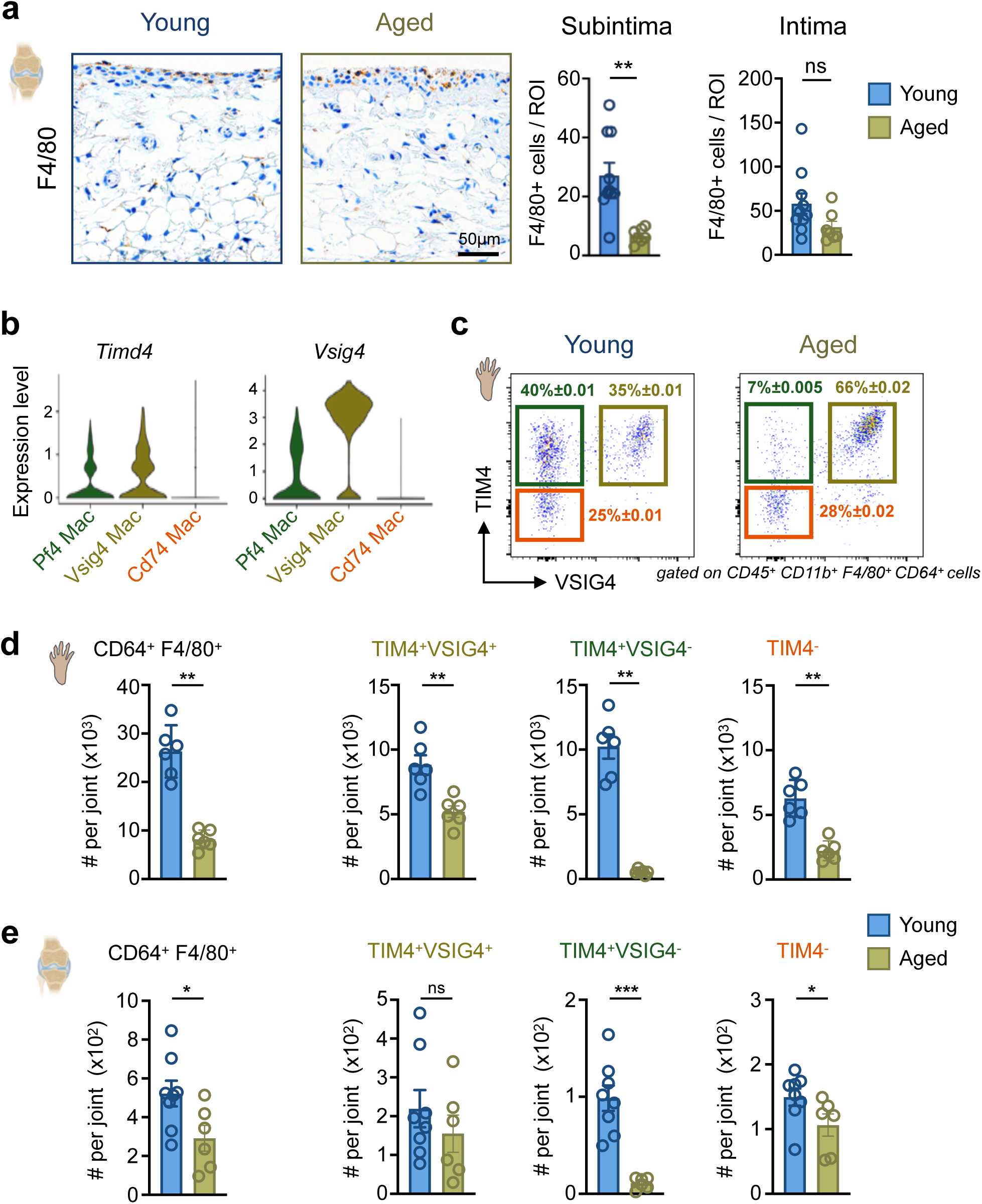
TIM4^+^VSIG4^−^ sublining macrophages are predominantly affected upon aging. **a,** Representative F4/80 immunohistochemistry in knee synovium from young and aged mice. Bar graphs on the right show quantification of F4/80^+^ cells in the synovial subintima and intima (*n* = 10 young and *n* = 7 aged mice). Scale bar, 50 µm. **b**, Violin plots showing *Timd4* and *Vsig4* expression across scRNA-seq-defined synovial macrophage subsets. **c**, Flow cytometry plots showing TIM4 and VSIG4 expression within CD64^+^F4/80^+^ synovial macrophages from young and aged ankle joints. **d**, Quantification of total CD64^+^F4/80^+^ and macrophage subsets segregated by TIM4 and VSIG4 protein expression in the ankle joints of young and aged mice (*n* = 6 per group). **e**, Enumeration of total CD64^+^F4/80^+^ macrophages and TIM4/VSIG4-defined macrophage subsets in the knee synovium of young (n = 8) and aged (n = 6) mice by flow cytometry. Data are mean ± s.e.m. Two-tailed Mann-Whitney *U* test. \**P* < 0.05, \*\**P* < 0.01, \*\*\**P* < 0.001; ns, not significant.

### Loss of TIM4^+^VSIG4^−^ sublining macrophages correlates with increased synovial fibrosis

To further assess the functional relevance of this age-reduced macrophage niche, we examined its relationship to published macrophage datasets and matrix-associated functions. Scoring against the Lyve1⁺ and Lyve1^−^ macrophage profiles described by Chakarov et al. across multiple tissues showed that resident synovial macrophages strongly overlap with a conserved Lyve1^+^ macrophage gene signature (Fig. 7a)^23^. This was supported by expression of transcript and protein markers, including *Mrc1* (encoding CD206) and *Lyve1* (Fig. 7b,c; Supplementary Table 6). LYVE1⁺ macrophages have been linked to tissue-supportive and matrix-regulatory functions, including collagen homeostasis^23–26^. Bulk RNA-seq of FACS-sorted TIM4⁺ synovial macrophages showed clear age-dependent separation and remodeling of the TIM4⁺ compartment (Extended Data Fig. 10a,b). Genes reduced with age mapped to the *Pf4* macrophage state when projected onto the single-cell dataset, consistent with the loss of the sublining component of the TIM4⁺ pool (Extended Data Fig. 10c). Gene set enrichment analysis further revealed reduced representation of pathways linked to extracellular matrix and collagen degradation in aged TIM4⁺ macrophages (Extended Data Fig. 10d). Pan-macrophage depletion in young CD64-DTR mice increased synovial collagen accumulation, recapitulating a fibrotic feature of the aged joint. In contrast, depletion of CX3CR1⁺ lining macrophages while sparing other synovial macrophages did not induce a comparable fibrotic phenotype (Fig. 7d; Extended Data Fig. 10e,f). Altogether, these findings link age-associated loss of TIM4⁺ sublining macrophages to reduced extracellular matrix homeostasis and support the idea that this macrophage niche contributes to limiting fibrotic remodeling in the joint.

**Fig. 7:**
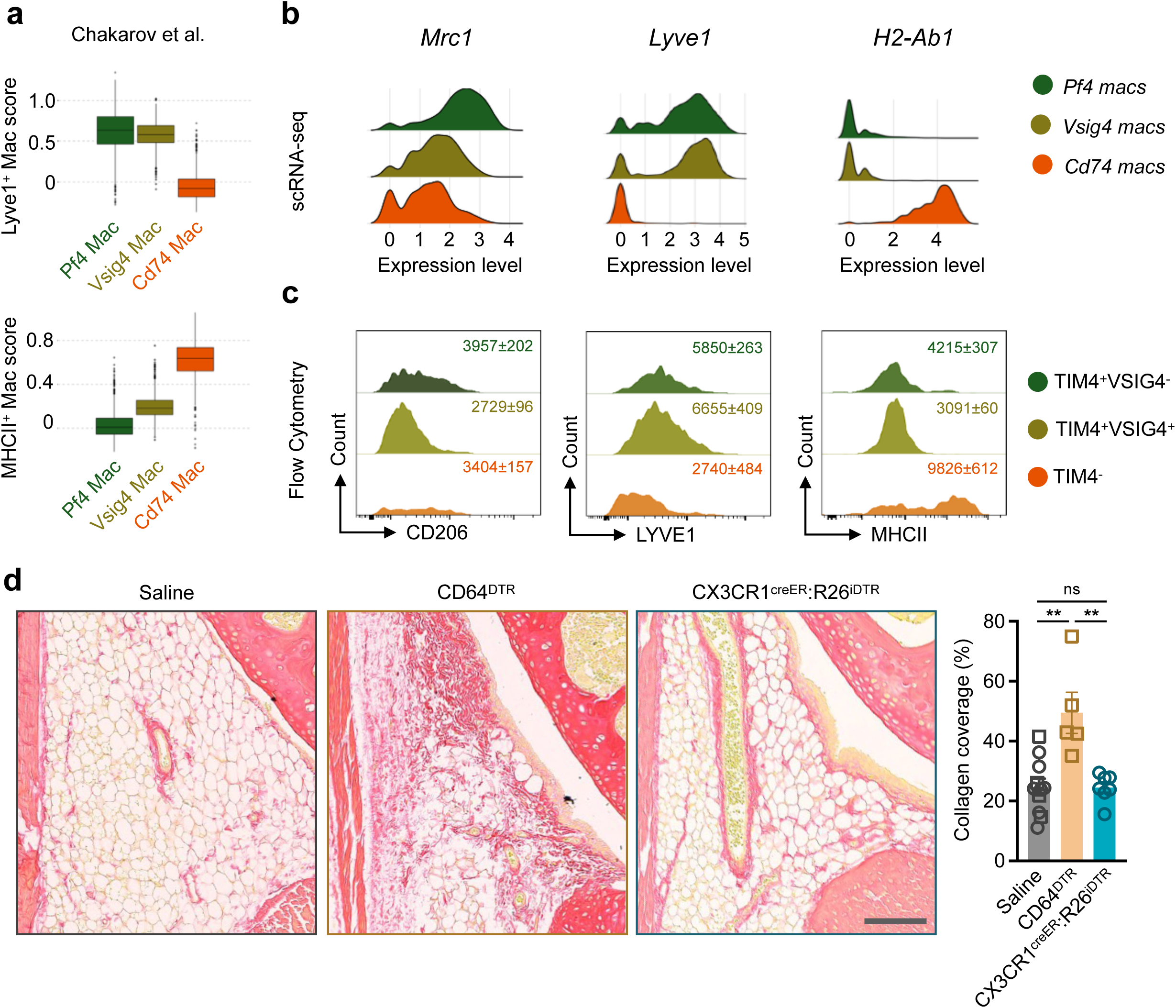
TIM4^+^VSIG4^−^ macrophages support extracellular matrix homeostasis and limit synovial fibrosis. **a,** Module scores for published LYVE1^+^ and MHC-II^+^ macrophage signatures from Chakarov et al. mapped across synovial macrophage subsets. **b,** Gene expression of *Mrc1, Lyve1* and *H2-Ab1* across *Pf4*^+^, *Vsig4*^+^ and *Cd74*^+^ macrophage subsets. **c,** Flow cytometry histograms showing CD206, LYVE1 and MHCII expression across macrophage subsets delineated by TIM4 and VSIG4 cell surface markers. **d,** Representative Sirius Red-stained knee synovium in saline-treated control mice (n = 11), diphtheria toxin-treated CD64-DTR mice (n = 5) and diphtheria toxin-treated CX3CR1creER-iDTR mice (n = 7). Bar graph on the right shows collagen coverage quantification. Saline treated animals includes littermate controls from both transgenic lines. Scale bar, 100 µm. Data are mean ± s.e.m. Kruskal-Wallis test. \*\**P* < 0.01; ns, not significant.

## Discussion

Aging is a major risk factor for joint disorders, yet the intrinsic cellular remodeling of the synovium during aging remains incompletely defined. In this study, we provide a single-cell transcriptomic atlas of the physiologically aging murine synovium, establishing a first reference map for defining the baseline cellular landscape of age-related joint vulnerability. Aged synovium is characterized by remodeling within the sublining stromal-myeloid niche, including fibroblast oxidative stress programs, sublining fibrosis, and selective attrition of resident fibroblast and macrophage populations. Notably, these changes occur in the setting of systemic low-grade inflammation and increased myelopoiesis, but without the hallmarks of inflammatory synovitis, suggesting that age-associated joint vulnerability may instead reflect the progressive decline and dysfunction of protective resident niches. Our identification of resident macrophage loss as a prominent feature of synovial aging places the joint within a growing body of evidence indicating that niche-specific attrition of TRMs is a conserved hallmark of aging across organs^27–29^. Indeed, embryonically seeded TRMs including Kupffer cells in the liver, alveolar macrophages in the lung, and macrophages in the skin decrease upon aging without contribution of bone marrow-derived cells^30–32^. In the skin, for example, loss of dermal macrophages disrupts vascular maintenance and promotes fibrosis and capillary rarefaction, highlighting the emergence of TRM-poor niches as a contributor to tissue fragility during aging^30^. Consistent with this, our data support the idea that specific tissue resident macrophage niches can decline during aging without adequate compensation by circulating cells despite aging myelopoiesis bias, potentially reflecting cumulative stress and the decay of microenvironmental survival cues.

A notable feature of the TIM4^+^VSIG4^−^ synovial subset is its transcriptional resemblance to LYVE1^+^ macrophages described across diverse tissues, which are increasingly linked to anti-fibrotic functions and ECM organization^23,24^. Within the joint, this concept is particularly compelling in light of recent high-dimensional mapping of human rheumatoid arthritis (RA) synovium showing that a perivascular network of LYVE1⁺ TRM is disrupted during active disease, and that restoration of this homeostatic macrophage architecture is a hallmark of clinical remission^26,33^. The loss of LYVE1⁺ macrophages during physiological murine aging supports the idea that age-related attrition of these cells contributes to joint vulnerability by weakening local homeostatic control, thereby increasing susceptibility to joint disease, including osteoarthritis and RA.

The concomitant decline of sublining fibroblasts further strengthens that aging destabilizes intercellular networks that are critical for joint homeostasis. Fibroblast loss and stress-associated reprogramming mirror findings in other mesenchymal tissues, such as the skin, where dermal aging is associated with cell senescence, matrix thinning and disruption of mechanical integrity^34–36^. The close spatial and functional interplay between fibroblasts and macrophages in the sublining niche suggests that their decline may be interdependent: macrophage dysfunction could impair fibroblast support and matrix turnover, while fibroblast senescence or altered trophic signaling may, in turn, compromise macrophage maintenance. Disentangling these causal relationships will be important for further exploring the exact mechanisms driving age-dependent loss of synovial tissue homeostasis.

In addition, although macrophage depletion experiments support an indirect functional link between macrophage loss and fibrosis, we currently lack tools to selectively perturb LYVE1^+^ interstitial macrophages in synovium. Future studies utilizing more specific genetic mouse models will be necessary to directly target synovial macrophages seeding the sublining tissue niche. Another limitation of the current study is the capture of discrete age points, providing snapshots rather than a continuous trajectory of cellular changes. Longitudinal and lineage-tracing studies are needed to determine the temporal order of fibroblast and macrophage attrition and to identify the intrinsic and extrinsic factors driving their progressive loss ^37–39^. Furthermore, the potential contributions of less abundant synovial populations, such as endothelial cells, lymphocytes, and mast cells, warrant further investigation, particularly as additional niche partners that may tune synovial homeostasis. Incorporating spatial transcriptomics and nuclei-based approaches may as well help to resolve the role of uncaptured synovial adipocytes across aging^40^.

In summary, we provide a high-resolution resource mapping physiological synovial aging and identifying niche-specific loss and dysfunction of resident fibroblast and macrophage populations as hallmarks of synovial remodeling with age. By linking the joint to emerging evidence of TRM attrition across organs, our study highlights maintenance of resident stromal-immune niches as a potential axis for preserving tissue integrity during aging. These findings further suggest that therapeutic strategies aimed at restoring macrophage niches or limiting fibroblast stress may represent promising avenues to mitigate age-associated joint decline.

## Methods

### Animals

C57BL/6J mice were purchased from Charles River Laboratories, including young (2 to 3 months) and naturally aged (18 to 24 months) cohorts. Cx3cr1^CreER^ (B6.129P2(Cg)-*Cx3cr1*^tm2.1(cre/ERT2)Litt/WganJ^, 021160) and R26^iDTR^ (C57BL/6-*Gt(ROSA)26Sor*^tm1(HBEGF)Awai^/J, 007900) were purchased from the Jackson Laboratory. CD64^DTR^ (*Fcgr1-IRES-EGFP-hDTR*) mice were provided by Sandrine Henri (CIML)^41^. To deplete synovial lining macrophages, Cx3cr1^CreER^ transgenic mice were crossed with R26^iDTR^ to generate Cx3cr1^CreER/+^;R26^iDTR/+^ animals. Genotyping was conducted according to the protocols provided on the Jackson Laboratory website. Mice were bred and housed in our animal facility at the Institute for Neurosciences of Montpellier under specific pathogen-free conditions with ad libitum access to food and water. All animal experiments were approved by the French Ministry of Higher Education and Research, the Languedoc-Roussillon Animal Research Ethics Committee, and the French Health Authorities, in compliance with European guidelines for animal testing.

### Cell preparation and isolation

For knee joint cell isolation, synovial tissues were dissected and minced into small pieces as described previously^42^. For ankle joints, following hindlimbs harvesting, meticulous dissection of intact individual bones (including the calcaneus, navicular, cuboid, and cuneiform bones) was carried out to mitigate unwanted tissue contamination and preserve the synovial tissue covering the articular recesses. Cell suspensions were obtained upon enzymatic digestion using a solution containing Collagenase type IV (1mg/ml, Gibco), DNase I (0.03mg/ml, Sigma Aldrich) in RPMI-1640 medium supplemented with 2% fetal bovine serum (FBS), for 60 minutes at 37°C on a shaker at 800 RPM. Supernatants from digested samples were filtered through a 35-µm cell strainer, washed in phosphate-buffered saline solution (PBS), centrifuged at 340g for 5 minutes and resuspended in flow cytometry buffer (PBS supplemented with 4% FBS) for subsequent flow cytometry staining. Peripheral blood was collected by retro-orbital bleeding using heparinized capillary tubes, and red blood cells were lysed using RBC lysis buffer (BioLegend). Cells were rinsed in PBS and pelleted by centrifugation prior to resuspension in PBS 4% FBS and flow cytometry staining.

### Single-cell RNA sequencing library preparation

Live single synovial cells from ankle joints were purified by FACS using a Cytek Aurora cell sorter in PBS containing 0.04% bovine serum albumin (BSA) and processed immediately for library preparation. Viable single-cell suspensions were loaded onto a Chromium Controller (10x Genomics) for droplet-based scRNA-seq using the Chromium Single Cell 3’ Library and Gel Bead Kit v3. The validation of the libraries was carried out using DNA quantification on the Fragment Analyzer (High Sensitivity NGS kit) as well as by qPCR (ROCHE Light Cycler 480) prior to sequencing using a NovaSeq 6000 (Illumina) at the MGX Platform of Montpellier (www.mgx.cnrs.fr).

### scRNA-seq data processing and analysis

Raw sequencing data were demultiplexed and converted to FASTQ files using Cell Ranger mkfastq (v7.1.0). Single-cell gene expression matrices were generated with the Cell Ranger pipeline (v7.0.1, mm10-2020-A) and analyzed using Seurat v4 in R^43^. Quality control was performed to exclude cells with fewer than 200 features, more than 6,000 features, or mitochondrial content exceeding 8%. Following this quality control, a total of 6,362 cells from the young pool and 7,058 cells from the aged pool were retained for subsequent analysis. Datasets were then merged, and normalization was performed using SCTransform with regression of cell cycle and mitochondrial gene effects. Clustering analysis was conducted with a resolution parameter of 0.45. Differentially expressed genes (DEGs) were identified using the FindAllMarkers function in Seurat. Gene Ontology (GO) enrichment analysis was performed using ClusterProfiler^44^. Module scores were calculated at the single-cell level using Seurat’s AddModuleScore function. The SASP score was computed using the SenMayo gene set described by Saul et al.^12^. Fibroblast annotation was further supported by scoring cells against published synovial fibroblast taxonomy signatures from Collins et al^10^. Macrophage subsets were similarly compared using published signatures from Culemann et al. and Chakarov et al^20,23^. To integrate bulk and single-cell data, significantly upregulated and downregulated genes identified by bulk RNA-seq were used as gene modules and projected onto the corresponding single-cell populations. Regulon activity was inferred with pySCENIC, cell-cell communication was analyzed with CellChat, and the dataset was visualized using the UCSC Cell Browser^45–48^.

### Bulk RNA-seq

Cells collected by FACS from several animals were pooled. Total RNA was extracted using Nucleospin RNA Plus XS (Macherey Nagel), according to manufacturer’s protocol. All RNAs were analyzed on an Agilent Fragment Analyzer. Libraries were prepared with the SMART-Seq Stranded kit (Takara Bio) and were sequenced with a SP50 paired end flow cell on a NovaSeq 6000 (Illumina). Base calling was made with Illumina bcl-convert. Sequences were aligned using HISAT2^49^ with mm39 reference genome after FastQC quality assessment. featureCounts and DESeq2 were used for downstream analysis^50,51^. To perform Gene Set Enrichment Analysis (GSEA), the list of expressed genes from the DESeq2 analysis was ranked using a combined metric of the fold change direction and statistical significance: -log10(P-value) × sign(log2FoldChange). The resulting pre-ranked gene list was exported and analyzed using the GSEA desktop application (Broad Institute and UC San Diego)^52,53^.

### Flow cytometry

For blood and bone marrow samples, cells were stained with fluorophore-conjugated antibodies for 30 minutes at 4°C in PBS 4% FBS buffer. Cells were incubated with anti-CD45 (30-F11, BioLegend), -CD11b (M1/70, BD Biosciences), -CD115 (AFS98, eBioscience), -Ly6C (HK1.4, BioLegend). Blood monocyte subsets were identified as CD45^+^ CD11b^+^ CD115^hi^ Ly6C^low/high^. Neutrophils were identified as CD45^+^ CD11b^+^ CD115^−^ Ly6G^+^. For synovial tissue macrophages, cells were incubated with anti-CD45 (30-F11, BioLegend), -CD11b (M1/70, BD Biosciences), -Ly6G (1A8, BioLegend), - Ly6C (HK1.4, BioLegend), -F4/80 (BM8, BioLegend), -CD64 (REA286, Miltenyi Biotec), -Tim-4 (RMT4-54, BioLegend), -MHCII (M5/114.15.2, BioLegend), -CD31 (MEC13.3, BioLegend), -Podoplanin (8.1.1, BioLegend), -PDGFRα (APA5, BioLegend), -CD90.2 (30-H12, BioLegend). When anti-biotin antibodies were used, BUV395-conjugated streptavidin (BD Biosciences) was added during a second incubation step. Synovial lining macrophages were identified as CD45^+^CD11b^+^F4/80^+^CD64^+^Ly6C^lo^ TIM4^+^ VSIG4^+^. Sublining macrophages were defined as CD45^+^CD11b^+^F4/80^+^CD64^+^Ly6C^lo^ TIM4^+^ VSIG4^−^. Monocytes in the synovium were identified as CD45^+^CD11b^+^ F4/80^lo^CD64^lo^Ly6C^+^ and neutrophils were CD45^+^CD11b^+^F4/80^lo^CD64^lo^Ly6G^+^. Synovial fibroblasts were defined as CD45^−^CD31^−^PDGFRα^+^Podoplanin^+^ and further assessed for Thy1 to discriminate between lining- (Thy1^−^) and sublining fibroblast (Thy1^+^) subsets, whereas endothelial cells were defined as CD45^−^CD31^+^. CellROX green staining was performed according to manufacturer’s protocol (Invitrogen, C10492). For in vivo synovial macrophage phagocytosis assay, pHrodo green *E. coli* bioparticles (20 µl) was injected peri-articularly in hind paws from young and aged mice two hours before tissue harvesting and processing. Absolute cell numbers were obtained with counting beads (Precision Count Beads™, BioLegend) and expressed per joint or ml of blood. Samples were pre-gated on single and viable cells using either eBioscience™ Fixable Viability Dye eFluor™ 780 or Sytox Blue. Cell sorting experiments were conducted using an Aurora CS cell sorter (Cytek®). Data acquisition was performed using a BD FACSymphony™ A3 Cell Analyzer (BD Biosciences) or a Cytek Aurora flow cytometer (Cytek®) and were analyzed using FlowJo software (v10.10.1).

### Ex vivo stimulation of synovial macrophages

FACS-sorted CD45^+^ CD11b^+^ F4/80^+^ CD64^+^ ankle joint macrophages from young and old mice were seeded into 96-well plates (20,000 cells per well) in the presence of either control medium (RPMI with 10% FBS and 1% penicillin/streptomycin) alone, or with 10 ng/mL of lipopolysaccharide (LPS) for 24 hours. Cell supernatants were collected and stored at -20°C.

### ELISA

Mouse TNFα (DY410, R&D Systems) and IL-6 (DY406, R&D Systems) concentrations in serum samples and cell culture supernatants were measured by ELISA according to the manufacturer’s instructions. For serum preparation, murine blood was collected, allowed to clot at room temperature for 30 min, and centrifuged at 2000 g for 15 min. The serum fraction was harvested and stored at −80°C until analysis.

### *In vivo* macrophage depletion experiments

Macrophage depletion experiments were performed using CD64^DTR^ mice and Cx3cr1^CreER/+^;R26^iDTR/+^. In Cx3cr1^CreER/+^;R26^iDTR/+^ mice, tamoxifen was administered intraperitoneally every two days for a total of three injections (200µL at 20mg/mL per injection) to induce recombination, and depletion was initiated one month later. In both models, diphtheria toxin (DT) was administered intraperitoneally on days 0, 2, and 5, whereas littermate controls received saline solution (NaCl 0.9%) following the same schedule. Mice were sacrificed on day 6 and tissues were collected for downstream analyses.

### Histology and immunohistochemistry

Mouse hind paws and knees were harvested and fixed for 48 hours in 4% paraformaldehyde (PFA) at room temperature. Specimens were washed, decalcified in 14% EDTA-PBS for three weeks with solution exchanges, and placed in 70% ethanol prior to paraffin embedding. Hematoxylin-eosin-saffron (HES) and Red Sirius staining were performed on tissue sections (3µm) at the Experimental Histology Network of Montpellier (RHEM; www.rhem.cnrs.fr) according to standard protocols. For FAP and F4/80 detection by immunohistochemistry (IHC), sections were deparaffinized, rehydrated and subjected to heat-induced antigen retrieval with pH6 buffer (Agilent, EnVision FLEX Target Retrieval Solution Low, #K800521-2) for 20 h at 60°C. IHC was performed on a VENTANA Discovery Ultra automated staining instrument (Ventana Medical Systems), using VENTANA reagents, according to the manufacturer’s instructions. After neutralization of endogenous peroxidase activity, sections were incubated with anti-FAP (Abcam, ab207178, clone EPR20021) or anti-F4/80 antibody (Invitrogen, MF48000, clone BM8) for 60 min at room temperature. Signal enhancement was performed using the Discovery HQ conjugated antibody anti-rabbit IgG (cat# 07017812001, Roche Diagnostics) then Discovery amplification anti-HQ HRP Multimer (cat# 06442544001) according to the manufacturer’s instructions. Slides were incubated with DAB (cat# 05266645001) then counterstained with hematoxylin II (cat# 790-2208) for 8 min, followed by Bluing reagent (cat# 760-2037) for 4 min. Slides were then dehydrated with Leica autostainer and coverslipped with Pertex mounting medium with CM6 coverslipper (Microm). Brightfield stained slides were digitized with a Hamamatsu NanoZoomer 2.0-HT scanner and images were analyzed using QuPath software (v0.2.0)^54^. For Red Sirius staining, a pixel classifier was trained outside the analyzed area, and synovial tissue was isolated to analyze fibrosis content. For FAP and F4/80 quantifications, DAB stain deconvolution was performed and a threshold on DAB was used to detect positive cells/pixels.

### Statistical methods

Statistical analyses were performed using GraphPad Prism (version 9.5.1; GraphPad Software). Data are shown as mean ± s.e.m. The sample size and statistical test used for each experiment are provided in the corresponding figure legends. Two-group comparisons were performed using two-sided Mann-Whitney tests. For comparisons involving more than two groups, a Kruskal-Wallis test with Dunn’s multiple-comparisons test or one-way ANOVA with Šídák’s multiple-comparisons test was used, as appropriate. P values < 0.05 were considered statistically significant.

## Data availability

All scRNA-seq and bulk RNA-seq raw data were deposited in GEO and are available using the SuperSeries accession number GSE299534. An interactive website for this scRNA-seq dataset is available at https://aging.btlt.fr. Other research data are available upon reasonable request.

## Code availability

The scRNA-seq and bulk RNA-seq data analysis was performed using scripts in R, which are available in the GitHub repository: olivierbortolotti/synovial-mouse-aging (https://github.com/olivierbortolotti/synovial-mouse-aging).

## Acknowledgements

This work was supported by grants from the French National Research Agency (ANR-21-CE14-0077-01 to G.C.). The funders had no role in study design, data collection and analysis, decision to publish or preparation of the manuscript. We acknowledge the “Réseau d’Histologie Expérimentale de Montpellier” - RHEM facility for histology techniques and expertise. RHEM facility is supported by REACT-EU (Recovery Assistance for Cohesion and the Territories of Europe), IBiSA, Ligue contre le cancer, the Occitanie / Pyrénées-Méditerranée and GIS FC3R whose funds are managed by Inserm. SCENIC analysis was performed on the HPC facilities of the National Network of Computing Resources of the Institut Français de Bioinformatique (IFB), funded by the Programme d’Investissements d’Avenir (PIA), grant Agence Nationale de la Recherche number ANR-11-INBS-0013

## Contributions

O.B., F.A. and G.C. conceived the project and designed the experiments. O.B. and G.C. performed the experiments and data analyses, with contributions from L.M., B.A-A., D.S., F.L., C.D., L.B., J.C., C.B-G. and S.S.. O.B. and G.C. created figures and performed statistical analyses, wrote the manuscript, and all authors edited and approved the final draft.

## List of Supplementary tables

Supplementary Table 1: Global scRNA-seq Cluster Markers

Supplementary Table 2: Age-Associated DEGs in the scRNA-seq Atlas

Supplementary Table 3: Fibroblast scRNA-seq cluster Markers

Supplementary Table 4: Bulk RNA-seq DEGs for Fibroblasts

Supplementary Table 5: Macrophage scRNA-seq cluster Markers

Supplementary Table 6: Bulk RNA-seq DEGs for TIM4+ Macrophages

## Notes

https://aging.btlt.fr

### Competing Interest Statement

The authors have declared no competing interest.

