## Extended Data Fig for "Single-cell atlas reveals age-related cellular shifts underlying fibrosis in murine synovium"

**Extended Data Fig. 1 : Cell processing workflow and quality control for scRNA-seq of mouse ankle synovium.**

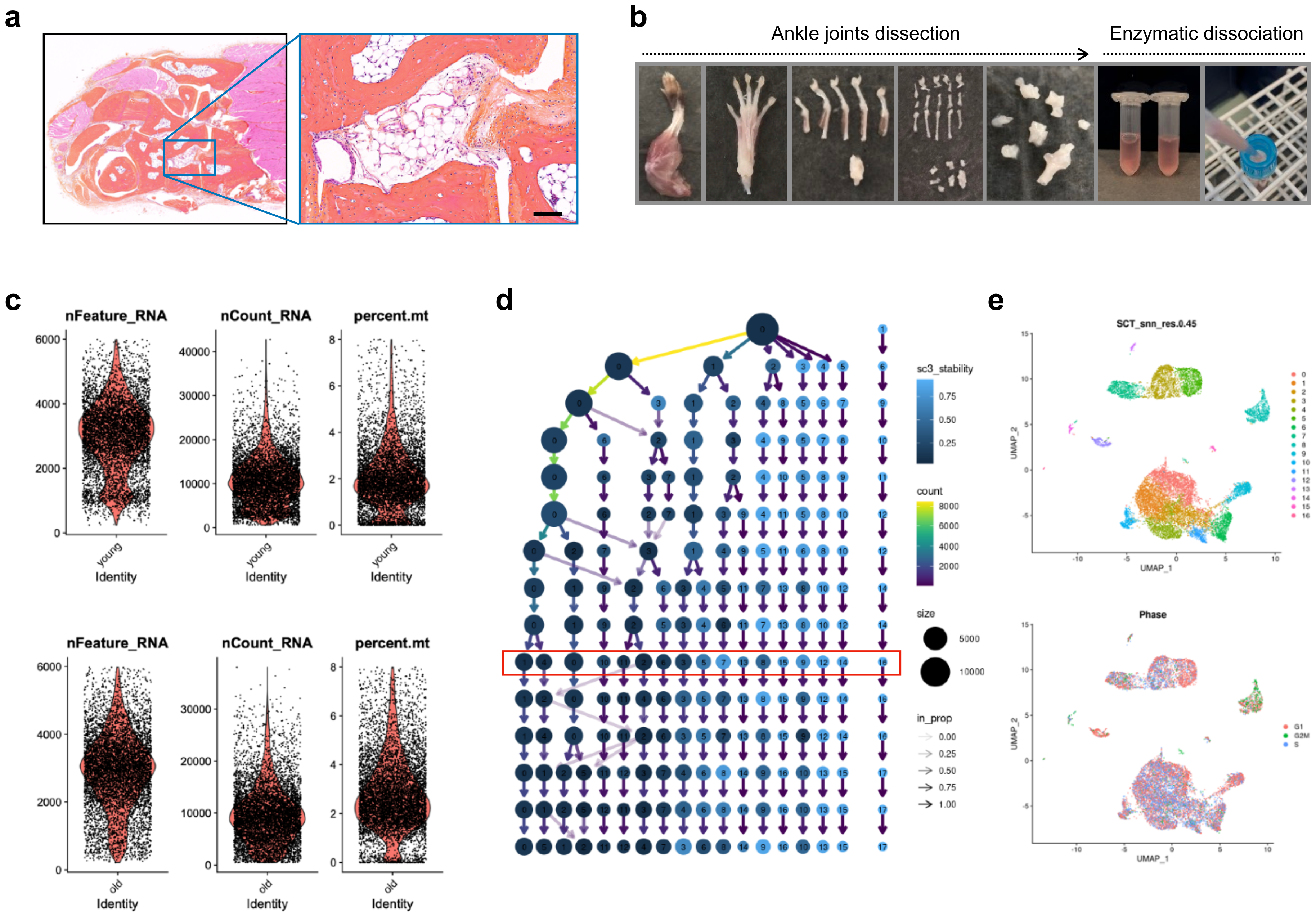

**Extended Data Fig. 2 : Canonical marker genes across mouse synovial cell populations.**

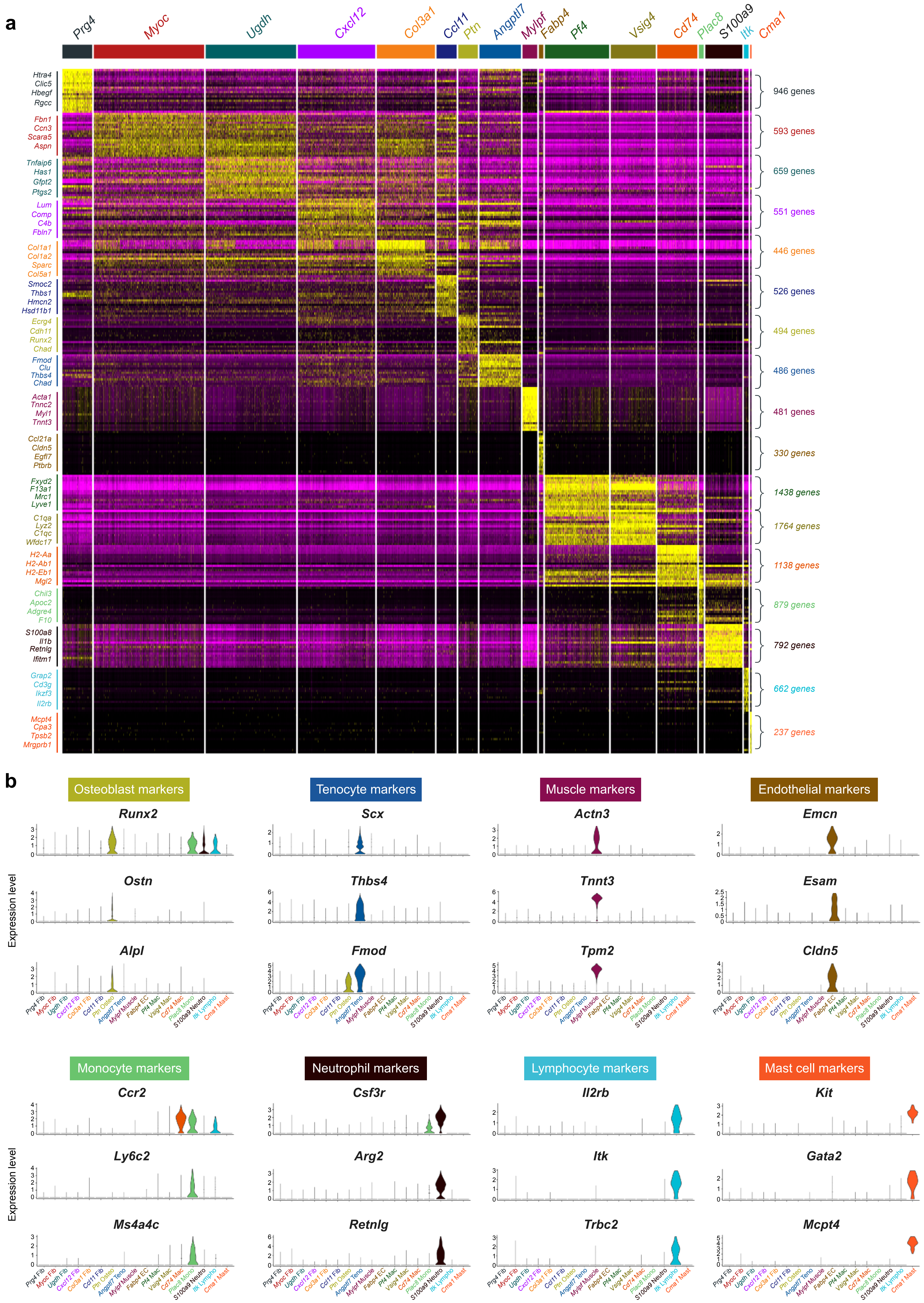

**Extended Data Fig. 3 : Synovial cell populations retain canonical marker expression with aging.**

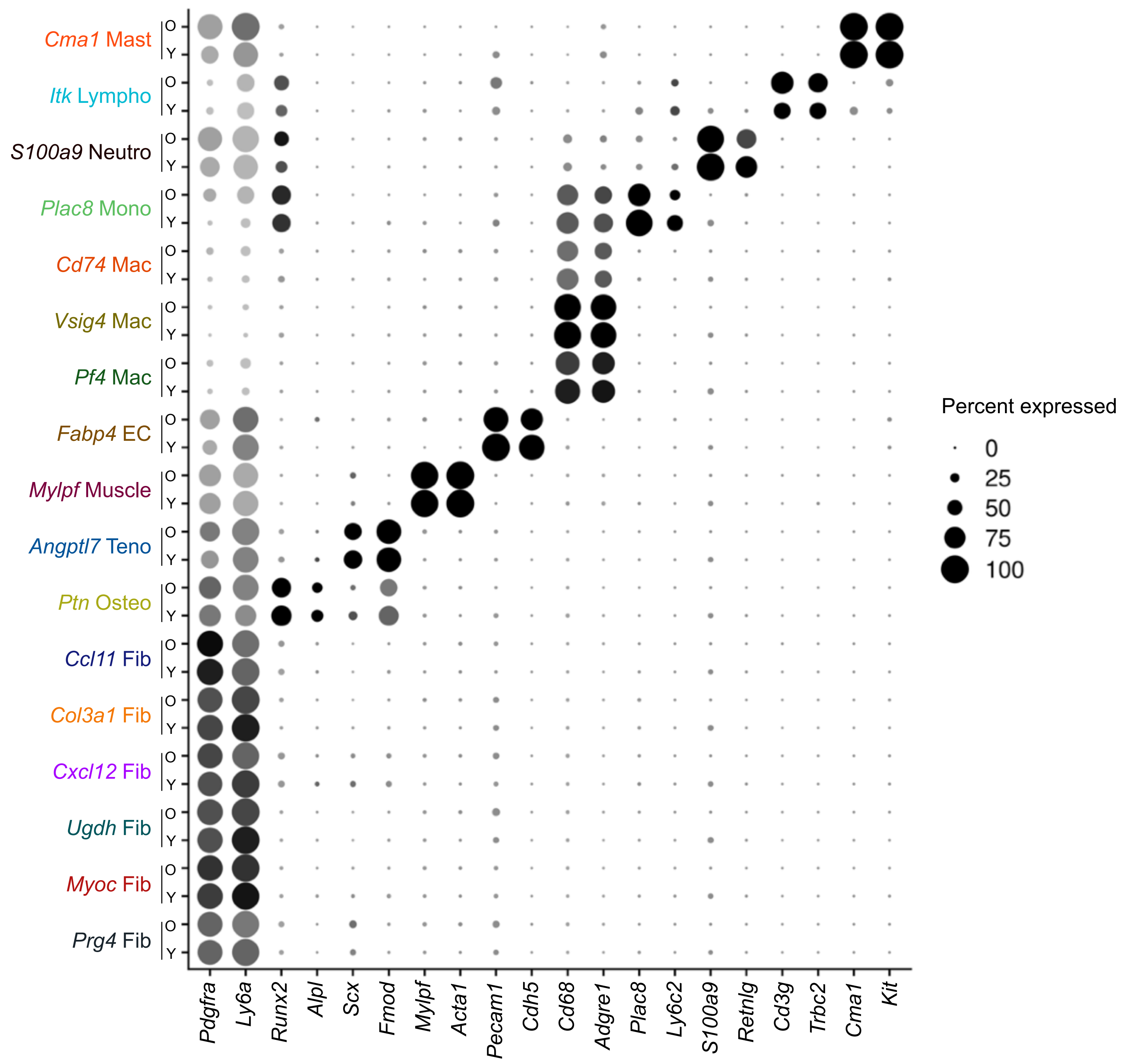

**Extended Data Fig. 4 : Functional annotation and comparison of synovial fibroblast clusters.**

**a**

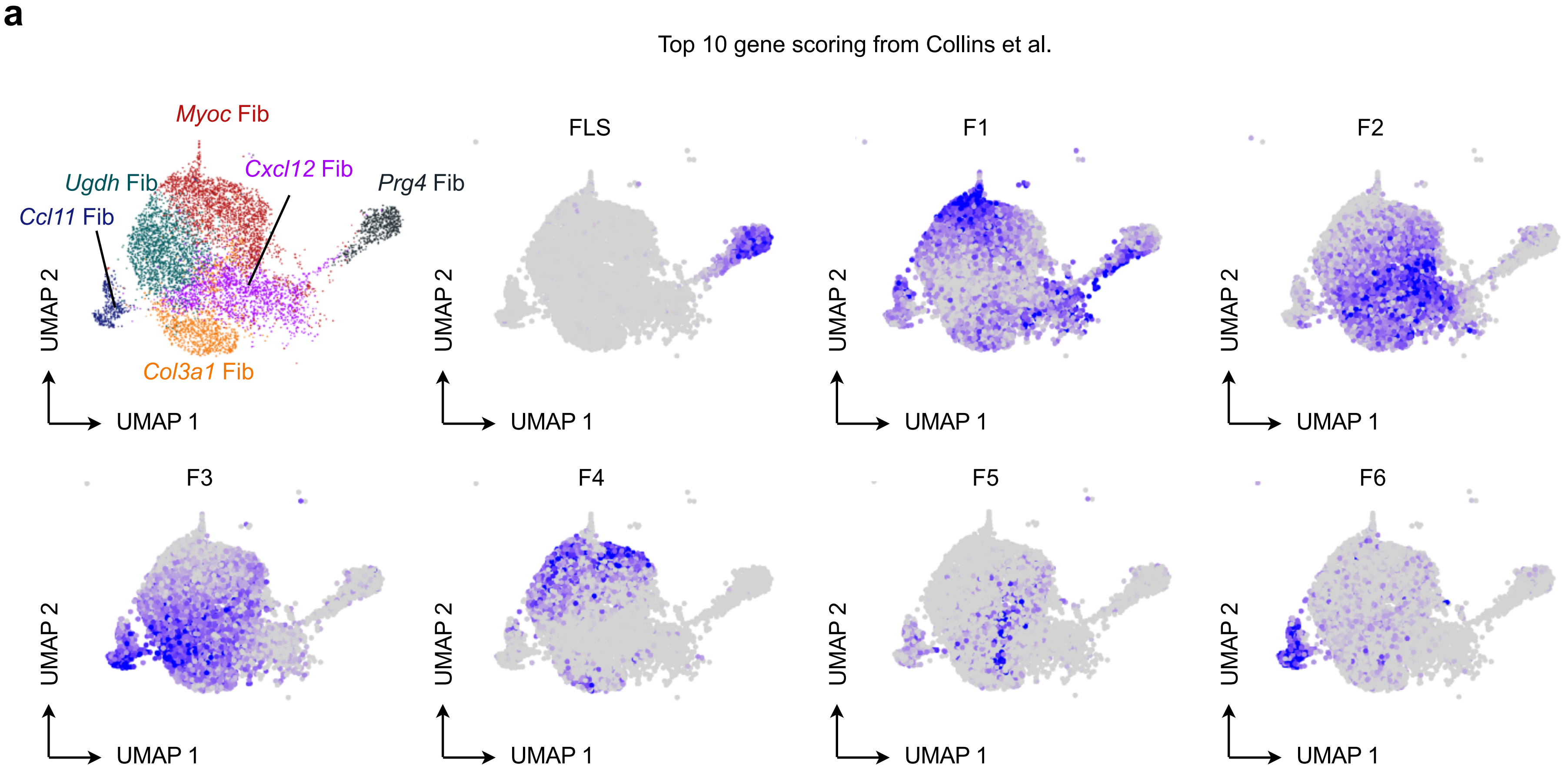


**b**

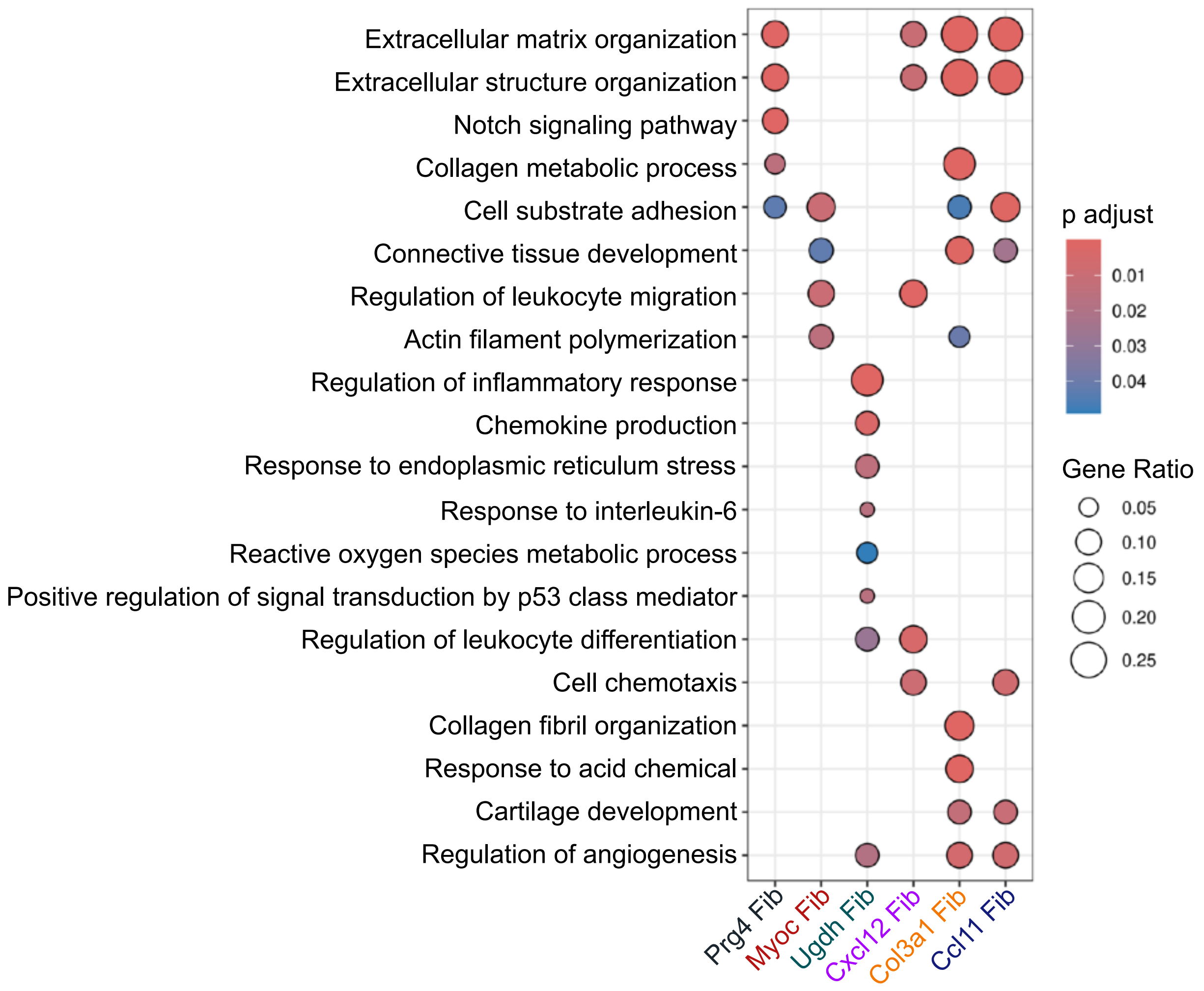

**Extended Data Fig. 5 : Intercellular communication networks and transcription factor regulon activity.**

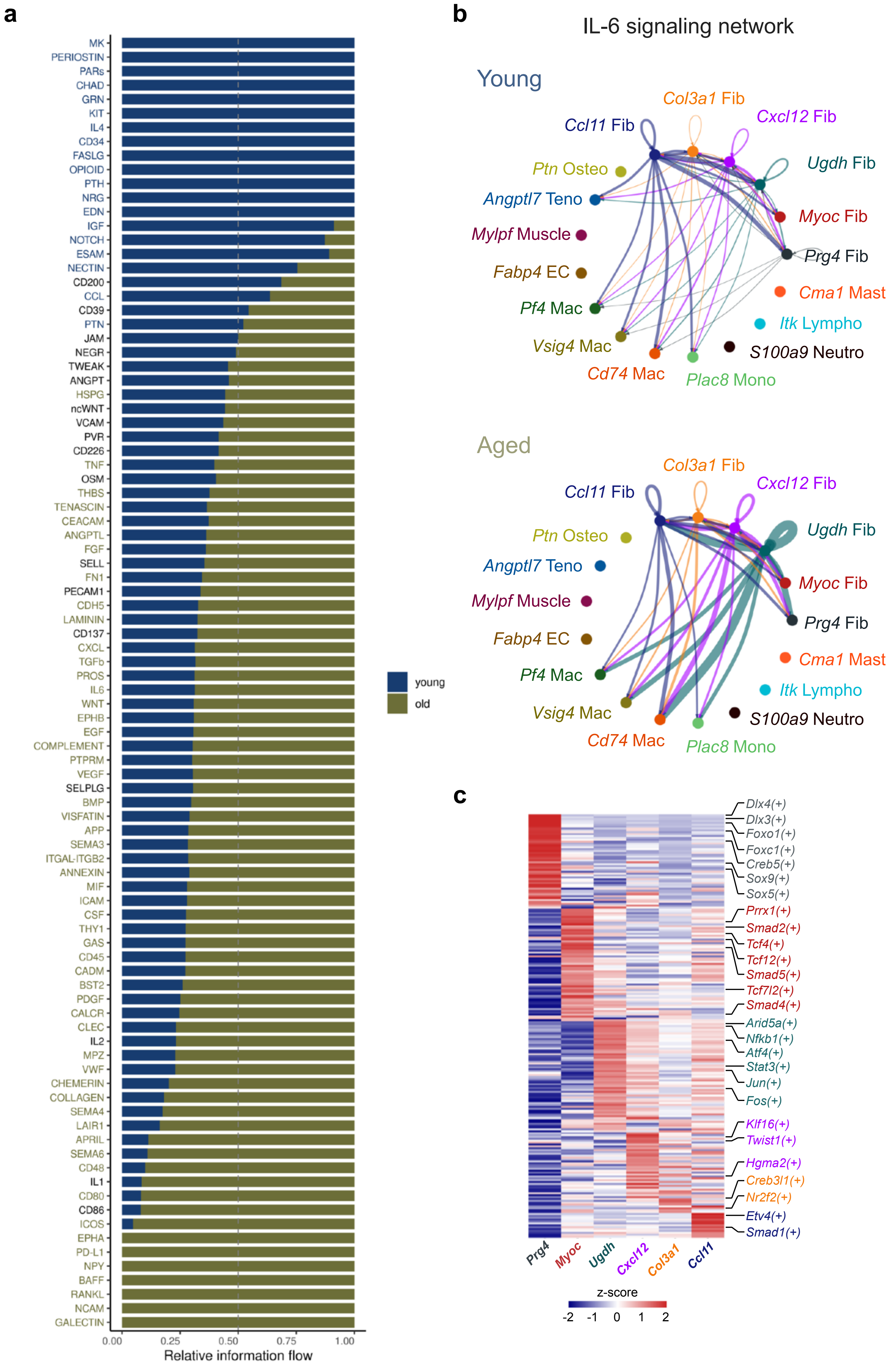

**Extended data Fig 6 : Synovial fibroblast decline coincides with ECM disruption and increased oxidative stress.**

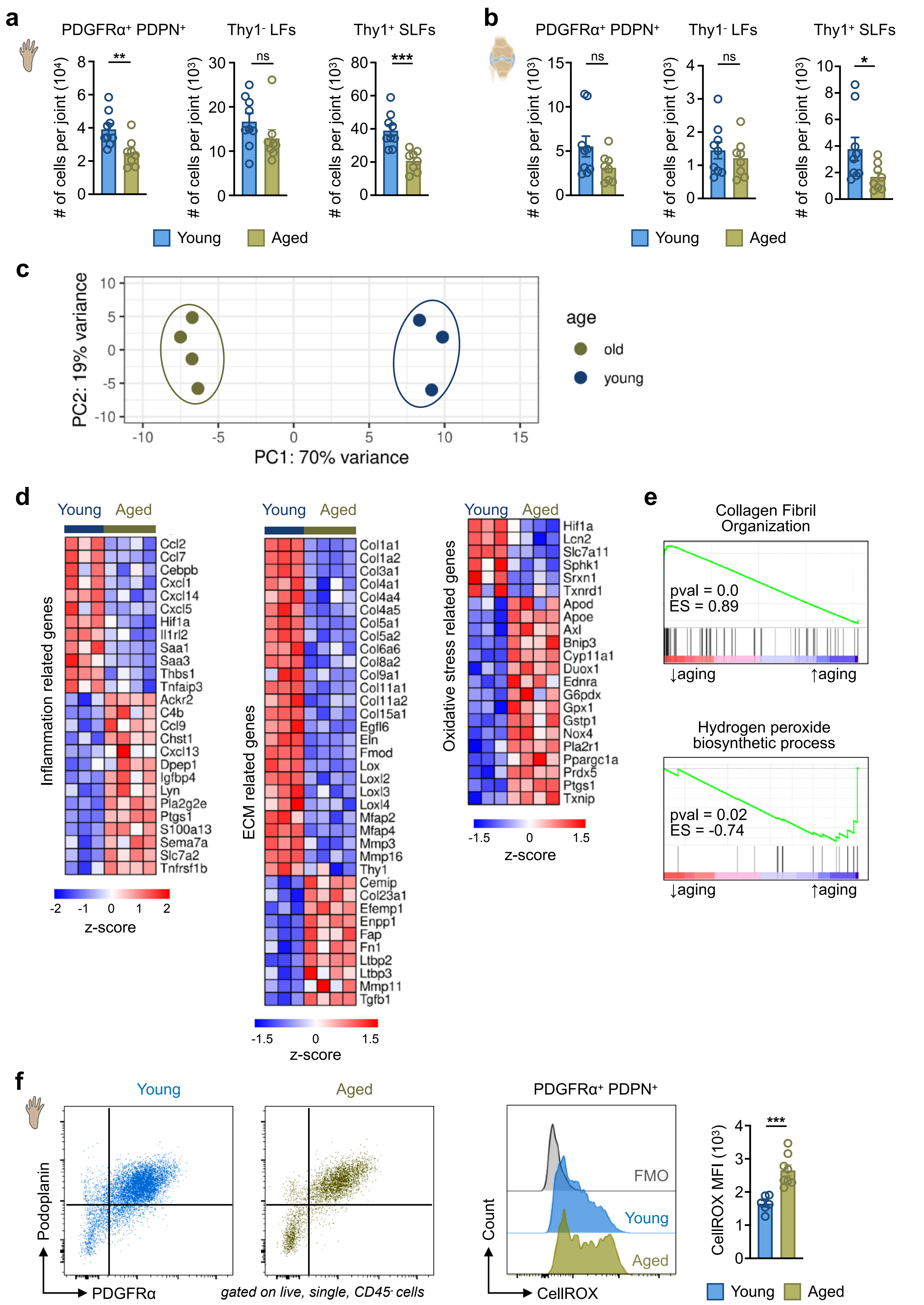

**Extended Data Fig. 7 : Systemic inflammatory and myeloid changes are conserved in aged female mice.**

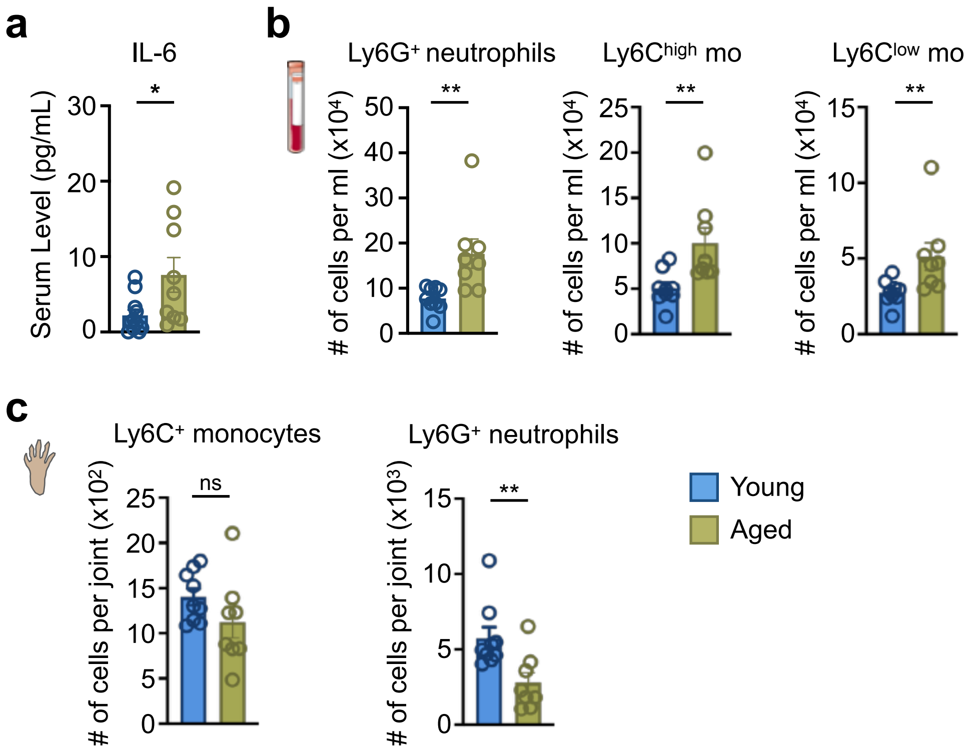

**Extended Data Fig. 8 : Functional annotation of synovial macrophage subsets.**

**a**

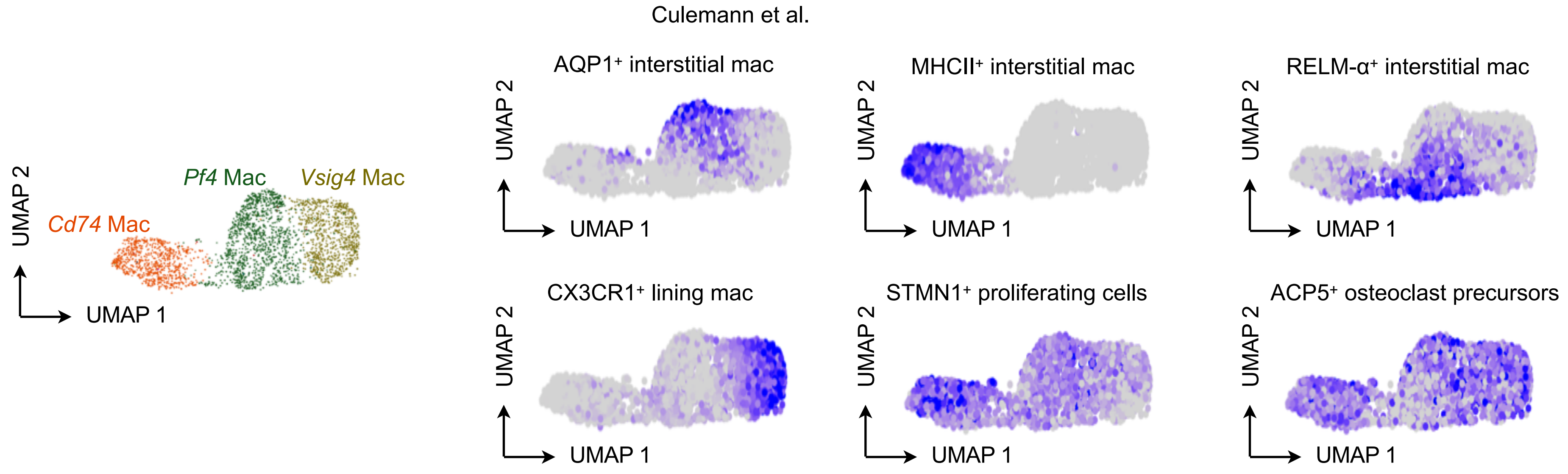


**b**

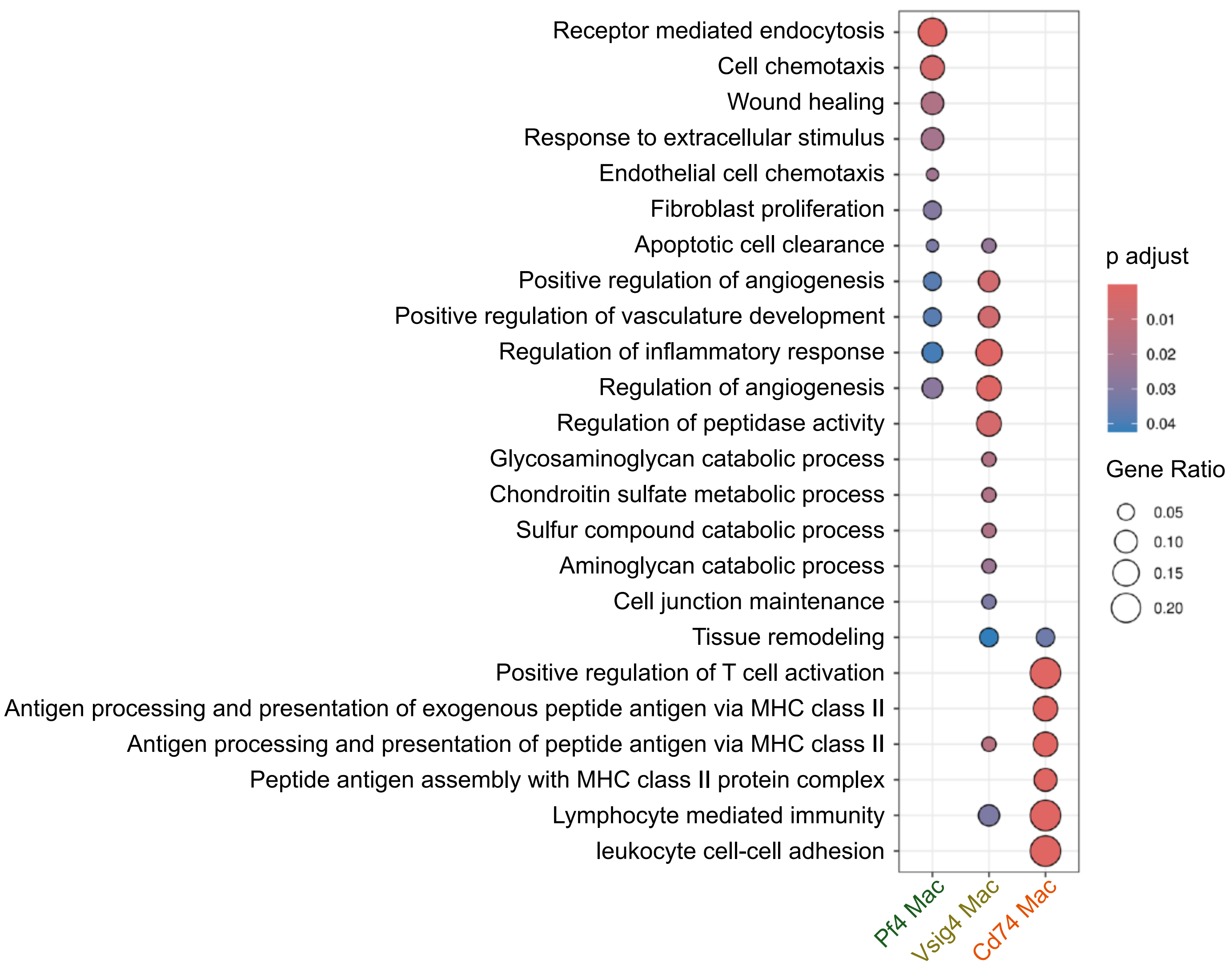

**Extended Data Fig. 9 : Age-related loss of TIM4⁺VSIG4⁻ macrophages is preserved in female mice.**

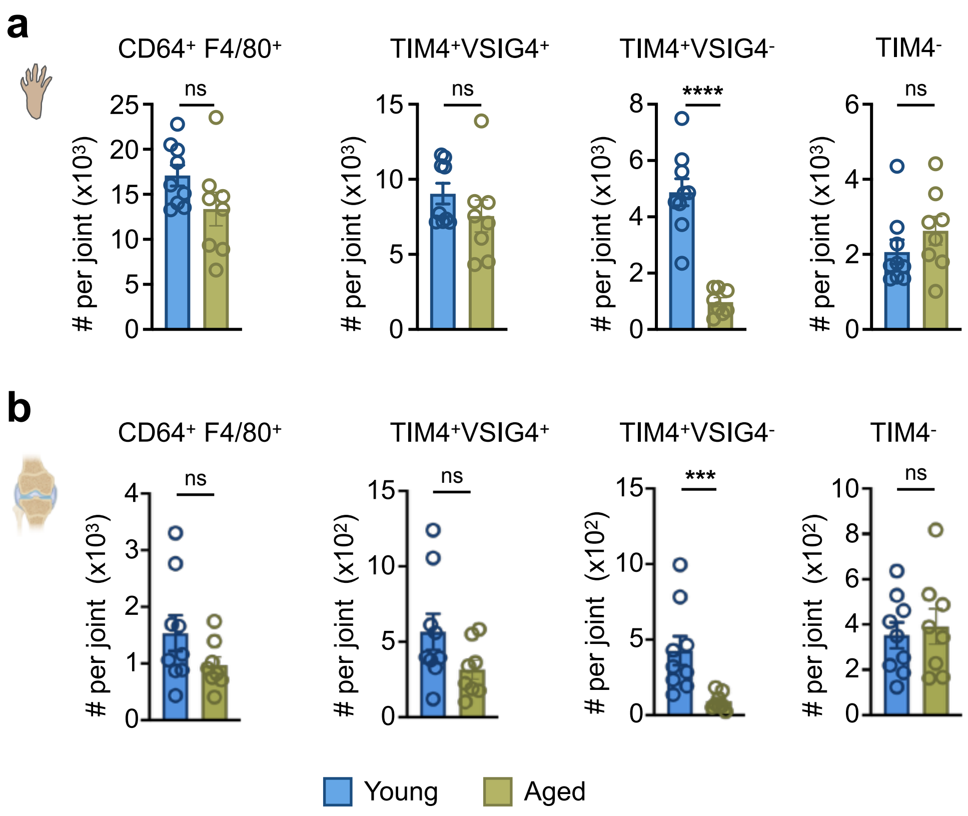

**Extended Data Fig. 10 : Loss of TIM4⁺VSIG4⁻ macrophages is associated with changes in ECM regulation.**

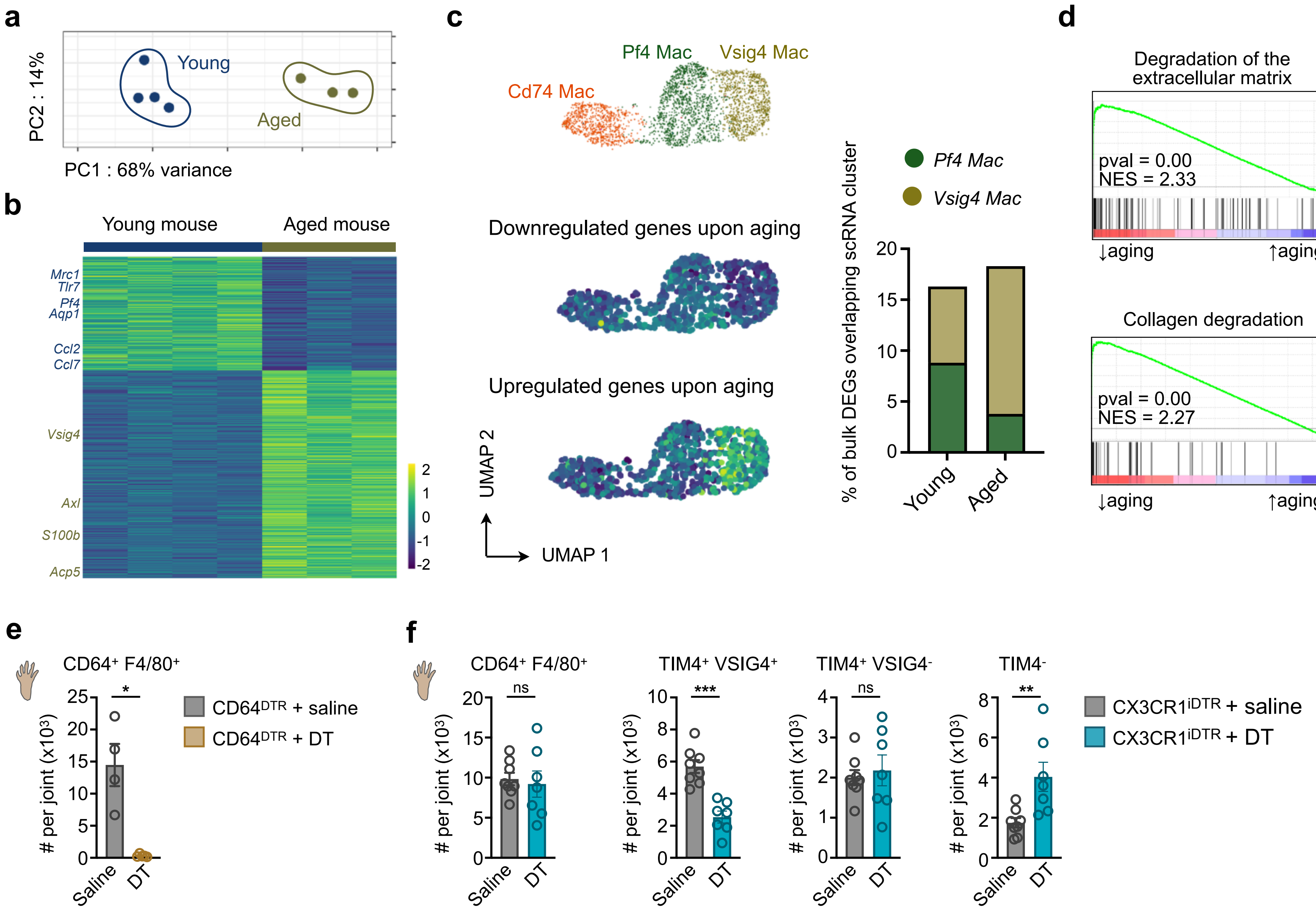
