## Extended Data Fig legend for "Single-cell atlas reveals age-related cellular shifts underlying fibrosis in murine synovium"

### Extended Data Figures

**Extended Data Fig. 1 : Cell processing workflow and quality control for scRNA-seq of mouse ankle synovium.**
**a,** Representative hematoxylin and eosin (H&E)-stained section of a mouse ankle joint showing the synovial compartment. Scale bar 100µm. **b,** Workflow for ankle joint dissection and enzymatic dissociation to generate single-cell suspensions. **c,** Quality control metrics for young (top) and aged (bottom) samples, showing the distributions of detected genes per cell (nFeature_RNA), total RNA counts per cell (nCount_RNA) and the percentage of mitochondrial transcripts (percent.mt). **d,** Clustering tree generated with Clustree across resolution parameters from 0 to 0.7 in increments of 0.05. The resolution used for downstream analyses (0.45) is highlighted in red. **e,** UMAP visualization of the scRNA-seq dataset at resolution 0.45 (top) and corresponding cell-cycle phase distribution after cell-cycle regression (bottom).

**Extended Data Fig. 2 : Canonical marker genes across mouse synovial cell populations.**
**a,** Heat map of the top 20 differentially expressed genes defining each synovial cell cluster identified by scRNA-seq. **b,** Violin plots showing expression of selected canonical marker genes for osteoblast, tenocyte, muscle, endothelial, monocyte, neutrophil, lymphocyte and mast cell populations.

**Extended Data Fig. 3 : Synovial cell populations retain canonical marker expression with aging.**
Dot plot showing expression of selected canonical marker genes across major synovial cell populations in young (Y) and aged (O) mice. Dot size indicates the percentage of cells expressing each gene.

**Extended Data Fig. 4 : Functional annotation and comparison of synovial fibroblast clusters.**
**a,** UMAP projections of published synovial fibroblast taxonomy scores from Collins et al. (FLS and F1-F6) across fibroblasts identified in this study. **b,** Gene Ontology enrichment analysis of biological processes associated with the top 50 DEGs defining each fibroblast cluster. Dot size indicates gene ratio and color indicates adjusted *P* value.

**Extended Data Fig. 5 : Intercellular communication networks and transcription factor regulon activity.**
**a,** Relative information flow for signaling pathways inferred by CellChat in young and aged synovium. **b,** Circle plots of inferred IL-6 signaling networks among synovial cell populations in young and aged mice. **c,** Heatmap showing SCENIC-inferred regulon activity for enriched transcription factors across fibroblast clusters.

**Extended Data Fig. 6 : Synovial fibroblast decline coincides with ECM disruption and increased oxidative stress.**
**a,** FACS-based quantification of total ($CD45^{-}CD31^{-}PDGFR\alpha^{+}PDPN^{+}$) fibroblasts, lining $THY1^{-}$ (LFs) and sublining $THY1^{+}$ fibroblast subsets in ankle joints from young (n = 9) and aged (n = 8) female mice. **b,** Enumeration of synovial fibroblast subsets from knee joints of female mice, shown as in a. (young n = 9 mice and aged n = 8 mice). **c,** Principal component analysis (PCA) of bulk RNA-seq of FACS-sorted synovial

fibroblasts isolated from young (n = 3) and aged (n = 4) mouse ankle joints. **d,** Heatmaps show selected differentially expressed genes related to inflammation, extracellular matrix and oxidative stress in young and aged synovial fibroblasts. **e,** GSEA plots comparing aged versus young synovial fibroblasts. **f,** Flow cytometry plots for PDGFRα$^+$PDPN$^+$ synovial fibroblasts and CellROX fluorescence histograms from young and aged mice are shown. Bar graphs on the right depict the mean fluorescence intensity (MFI) quantification of fibroblasts from young ($n$ = 6) and aged ($n$ = 8) animals. Data are mean ± s.e.m. Two-tailed Mann-Whitney $U$ test. *$P$ < 0.05, **$P$ < 0.01, ***$P$ < 0.001; ns, not significant.

**Extended Data Fig. 7 : Systemic inflammatory and myeloid changes are conserved in aged female mice.**
**a,** Serum IL-6 levels in young ($n$ = 11) and aged ($n$ = 9) female mice. **b,** FACS enumeration of Ly6C$^{high}$ monocytes, Ly6C$^{low}$ monocytes and Ly6G$^+$ neutrophils per ml of peripheral blood in young (n = 9) and aged (n = 8) female mice. **c,** Quantification of F4/80$^{low}$ Ly6C$^+$ monocytes and Ly6G$^+$ neutrophils in the ankle joints of young (n = 9) and aged (n = 8) female mice. Data are mean ± s.e.m. Two-tailed Mann-Whitney $U$ test. *$P$ < 0.05, **$P$ < 0.01; ns, not significant.

**Extended Data Fig. 8 : Functional annotation of synovial macrophage subsets.**
**a,** UMAP of synovial macrophages colored by cluster identity (*Pf4*+, *Vsig4*$^+$ and *Cd74*$^+$ subsets) and overlaid with module scores for published macrophage signatures from Culemann et al., including AQP1$^+$ interstitial, MHCII$^+$ interstitial, RELM-α$^+$ interstitial, CX3CR1$^+$ lining macrophages, STMN1$^+$ proliferating cells and ACP5$^+$ osteoclast precursors. **b,** GO enrichment analysis of the top 50 marker genes defining the *Pf4*$^+$, *Vsig4*$^+$ and *Cd74*$^+$ macrophage subsets. Dot size indicates gene ratio and color indicates adjusted $P$ value.

**Extended Data Fig. 9 : Age-related loss of TIM4$^+$VSIG4$^-$ macrophages is preserved in female mice.**
**a,b**, Flow cytometric quantification of total CD64$^+$F4/80$^+$ macrophages and TIM4/VSIG4-defined macrophage subsets per ankle (a) and knee joints (b) in young (n = 9) and aged (n = 8) female mice. Data are mean ± s.e.m. Two-tailed Mann-Whitney $U$ test. ***$P$ < 0.001, ****$P$ < 0.0001; ns, not significant.

**Extended Data Fig. 10 : Loss of TIM4$^+$VSIG4$^-$ macrophages is associated with changes in ECM regulation.**
**a,** Principal component analysis (PCA) of bulk RNA-seq from FACS-sorted TIM4$^+$ synovial macrophages isolated from young ($n$ = 4) and aged ($n$ = 3) male mice. **b,** Heatmap of differentially expressed genes between young and aged TIM4$^+$ synovial macrophages. **c,** UMAP projections of synovial macrophages scored for genes downregulated or upregulated with age in the bulk TIM4$^+$ macrophage dataset, with bar plot showing overlap of bulk differentially expressed genes with *Pf4* and *Vsig4* macrophage cluster signatures. **d,** GSEA plots comparing aged versus young TIM4$^+$ synovial macrophages, highlighting pathways related to extracellular matrix degradation and collagen degradation. **e,** Quantification of total CD64$^+$F4/80$^+$ macrophages per ankle joints in CD64$^{DTR}$ mice treated with saline (n = 4) or DT (n = 5). **f,** Flow cytometry quantification of total CD64$^+$F4/80$^+$ macrophages and subsets delineated by TIM4 and VSIG4 surface markers in ankle joints from CX3CR1-

$Cre^{ER}$;$R26^{iDTR}$ mice treated with saline (n = 8) or DT (n = 7). Data are mean ± s.e.m. Two-tailed Mann-Whitney *U* test. *$P < 0.05$, **$P < 0.01$, ***$P < 0.001$ ; ns, not significant.
